# The rapid regenerative response of a model sea anemone species *Exaiptasia pallida* is characterised by tissue plasticity and highly coordinated cell communication

**DOI:** 10.1101/644732

**Authors:** Chloé A. van der Burg, Ana Pavasovic, Edward K. Gilding, Elise S. Pelzer, Joachim M. Surm, Hayden L. Smith, Terence P. Walsh, Peter J. Prentis

## Abstract

Regeneration of a limb or tissue can be achieved through multiple different pathways and mechanisms. The sea anemone *Exaiptasia pallida* has been observed to have excellent regenerative proficiency but this has not yet been described transcriptionally. In this study we examined the genetic expression changes during a regenerative timecourse and report key genes involved in regeneration and wound healing. We found that the major response was an early upregulation of genes involved in cellular movement and cell communication, which likely contribute to a high level of tissue plasticity resulting in the rapid regeneration response observed in this species. We find the immune system is only transcriptionally active in the first eight hours post-amputation and conclude, in accordance with previous literature, that the immune system and regeneration have an inverse relationship. Fifty-nine genes (3.8% of total) differentially expressed during regeneration were identified as having no orthologues in other species, indicating that regeneration in *E. pallida* may rely on the activation of species-specific novel genes. Additionally, taxonomically-restricted novel genes, including species-specific novels, and highly conserved genes were identified throughout the regenerative timecourse, showing that both may work in concert to achieve complete regeneration. We conclude that *E. pallida* behaves similarly to other anemone species such as *Nematostella vectensis* and *Calliactis polypus* but with some notable novel differences.

## Introduction

Regeneration is defined as the recapitulation of development and is distinct from growth. All multicellular organisms must repair and regenerate tissue to some degree, in order to survive following injury or in context of daily cellular replacement. The ability to regenerate large portions of adult tissue, such as limbs or organs, is not universal among animal taxa, with even closely related species unable to regenerate to the same extent (Alvarado and Tsonis, 2006; Brockes et al. 2001; Liu et al. 2013; Poss, 2010; Tanaka and Reddien, 2011; Tiozzo and Copley, 2015). In general, the process of regenerating a tissue is as follows: first, the wound is closed and repaired in a process molecularly distinct from tissue regeneration. Then, precursor cells are mobilized for the formation of new tissue. Finally, the complete regenerated structure(s) is formed from the new tissue. Each of these steps may be achieved in ways unique to any one particular taxon or even be specific to a species (Alvarado and Tsonis, 2006; Brockes and Kumar, 2008; Garza-Garcia et al. 2010; Gurtner et al. 2008; Poss, 2010; Tanaka and Reddien, 2011; Tiozzo and Copley, 2015). Wound closure may proceed with or without cell proliferation, and new tissue may be formed from stem cells or through the trans- or de-differentiation of somatic cells (Tanaka and Reddien 2011).

A key pathway interaction that is often investigated is the role the immune system plays in regeneration. It has been hypothesised that the immune system may play an inhibitory role and typically, the more complex an organism’s immune system the less proficient the regenerative ability of that organism (Eming et al. 2009; Peiris et al. 2014). However, considering both regenerative and immune genes are typically activated early and with overlapping time frames following injury to an organism (Godwin and Brockes 2006; Eming et al. 2009; Peiris et al. 2014), it is plausible to hypothesise that some genes or pathways maybe shared or co-opted for both processes. Mixed results have been reported in the correlation of the two processes, with positive and negative influences found. For example, the immune system suppresses *Xenopus laevis* tadpole tail regeneration, with immunosuppressants restoring regeneration during the refractory period (Fukazawa et al. 2009). Conversely, in the coral *Acropora aspera*, the upregulation of the expression of putative immune system components and phenoloxidase activity was observed during regeneration post-injury (van de Water et al. 2015). Similarly, septic injury in the planarian *Schmidtea mediterranea* and the fresh water polyp *Hydra vulgaris* induced similar classes of genes involved in regeneration, at the same time the animals responded to immune challenge (Altincicek and Vilcinskas 2008).

Cnidaria is a phylum consisting primarily of marine animals, including sea anemones, hydroids, jellyfish and coral. Cnidarians have featured prominently among regeneration studies, due to their ease of use in a lab and their position in the phylogenetic tree as sister-phylum to Bilateria. One of the earliest animal model species is found in this phylum - the freshwater polyp *Hydra*, which has been a model organism for regeneration since 1744 (Browne, 1909; Gierer et al. 1972; Trembley, 1744). The *Hydra* and the more recent model organism the starlet sea anemone *Nematostella vectensis* have been used extensively as regeneration model organisms (Amiel et al. 2015; Bosch, 2007; DuBuc et al. 2014; Holstein et al. 2003; Passamaneck and Martindale, 2012; Petersen et al. 2015; Schaffer et al. 2016). Studies show that many genes, including genes involved in axis formation, tissue patterning and neural development, are present in even these morphologically simple animals. These were genes once thought to uniquely typify and shape the development of more complex animals, but may be the key to initiating regeneration through recapitulating development and reformation of tissue post injury (Babonis and Martindale, 2017; Forêt et al. 2010; Miller et al. 2005; Putnam et al. 2007; Sinigaglia et al. 2013).

Genome sequencing of *Exaiptasia pallida* (Baumgarten et al. 2015) has promoted its emergence as an important model species (Bucher et al. 2016; Oakley et al. 2016; Poole et al. 2016). Regeneration studies in *Exaiptasia* (Cnidaria: Actiniaria: Metridioidea), recently renamed as ‘*Exaiptasia*’ from ‘*Aiptasia*’ (Grajales and Rodríguez 2016) species are limited to the morphological changes that occur as *E. pallida* regenerates the tentacles and oral disc following transverse amputation (Singer and Palmer 1969; Singer 1971, 1974). Regenerating primary, secondary, tertiary and quaternary tentacles will emerge in a predetermined order around the oral disc and are distinguishable by size within a few days. Regeneration is able to proceed in *E. pallida* when DNA synthesis is blocked and collagen synthesis in the mesoglea increases substantially during regeneration (Singer 1974). *Exaiptasia pallida*, like many other cnidarians, can regenerate whole animals from small amputated parts and can reproduce both sexually and asexually via pedal laceration, making it amenable to clonal propagation in the lab (Clayton, Jr. 1985; Grawunder et al. 2015).

Here we describe the first transcriptomic study on *E. pallida* characterising the regenerative response to transverse amputation (head amputation). We generated multiple replicated datasets across seven timepoints to understand and delineate patterns of gene expression. We observe transcripts associated with cellular movement, signalling and response to be large components of the regeneration response. A substantial portion of novel differentially expressed genes is species-specific and a large portion is actiniarian-specific.

## Materials and Methods

### Ethics Statement

This project did not require animal ethics approval. Sample collection for sea anemones collected in Australia (all sea anemone datasets generated here, except for *Diadumene lineata*) was authorised under the Fisheries Act 1994 (General Fisheries Permit), permit number: 166312. Sample collection for *Diadumene lineata*, which was collected in New Zealand, was authorised under Ministry for Primary Industries Special Permit (632).

### Animal acquisition and visual observations

*Exaiptasia pallida* animals (n = 200) were collected from the Great Barrier Reef Marine Pty Ltd, Brisbane, QLD, Australia. Animals were kept in a holding tank at QUT until required for experimental use. To understand the stages of regeneration, animals were dissected transversely through the body column (perpendicular to the oral-aboral axis, as indicated in Figure 1), and then allowed to regenerate for several days. Visual observations of the animals were taken over several days and weeks before performing the time course assay. Five individuals were observed post-head amputation over at least 24 hours, up to 96 hours, using a Dinolite microscopic camera (DC20 Version:1.4.3). A representative video of this process is presented here (Supplementary Video File 1). To produce this video, four separate 23-hour films were taken over four consecutive days (the fourth video was slightly shorter at 19 hours of real time film) and were combined into one video, which shows 88 hours of total film occurring over 93 hours. A few hours were not recorded between separate videos. One second of this film represents about 20 minutes of real-time.

**Fig. 1.**
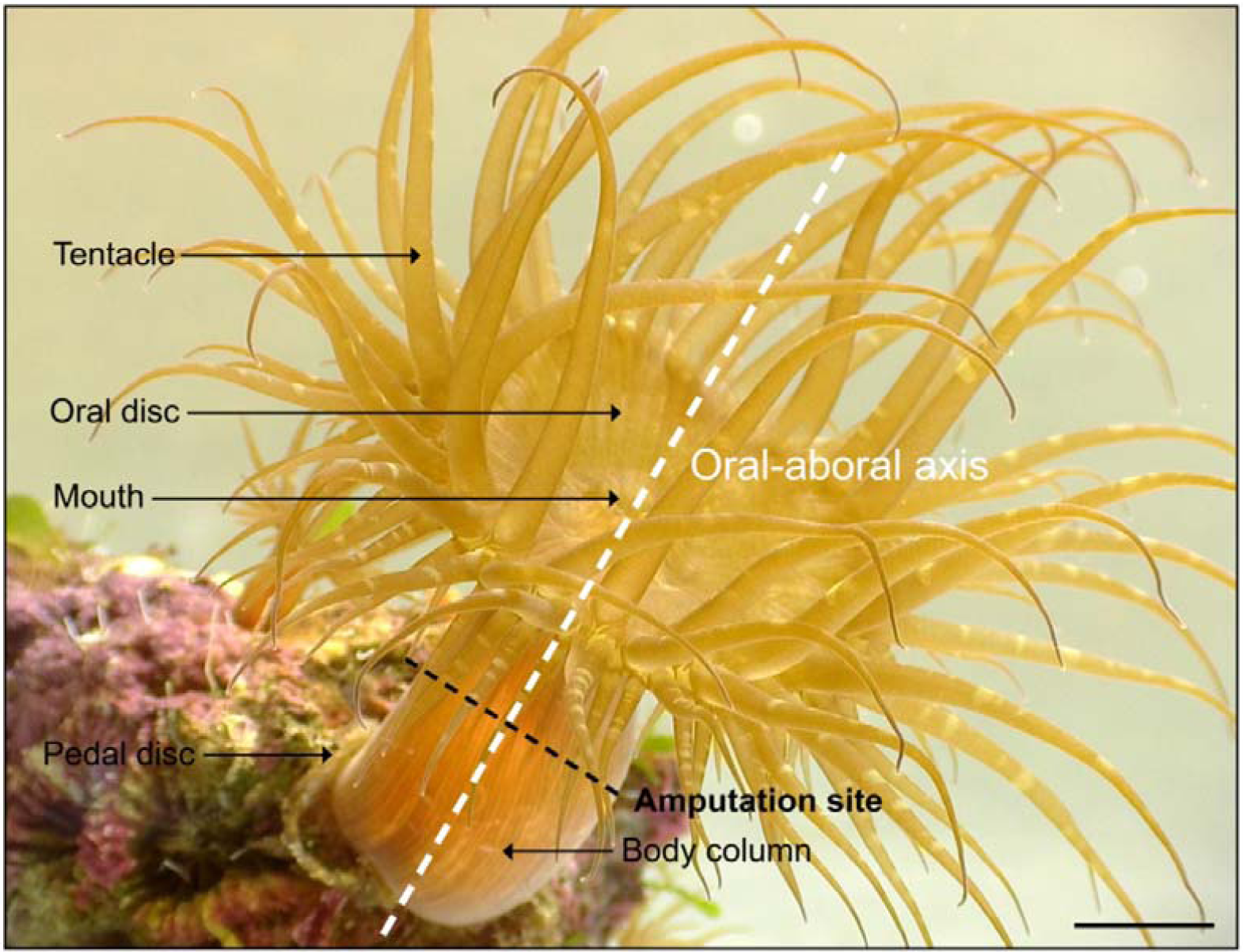
Diagram showing body layout, major axis and amputation site of *Exaiptasia pallida.* The image shows an adult individual attached via its pedal disc to a piece of live rock. The anemone is in the ‘extended relaxed’ position; its body column is extended upwards and elongated, and the tentacles are displayed openly. Scale bar = approx. 1 cm.

### Regeneration timecourse experiment

Animals (n = 21 per tank) were placed in three replicate tanks and maintained for one week with one feeding only and daily water changes, to allow animals to acclimate and attach to the tiles. Artificial sea water was maintained at between 35-37 ppt salinity and room temperature 20-21.5°C. Tiles were labelled with pencil to track replicates and ensure that an even representation of different sized animals was placed in each treatment group. Dissections were performed with surgical dissecting scissors through the body column, perpendicular to the oral-aboral axis (Figure 1). The dissected upper portions of the animals (the oral part consisting of part of the body column with the oral disc) were removed. The lower portion of the body (the aboral part of the individual which consists of the remaining column and pedal disc) remained in the replicate tanks until samples were taken at each of the seven time points (0 h, 45 m, 2.5 h, 8 h, 20 h, 48 h and 72 h). The 0 h sample (control) was taken immediately after head amputation, in order to capture baseline transcriptional activity. Singular samples consisted of three individual bodies combined from three tanks, which were immediately frozen in liquid nitrogen and then placed in the −80°C freezer for storage. Three replicates of each sample for each time point were generated.

### RNA-seq and library preparation

One sample for RNA-seq was produced by pooling three animals (described above), homogenising and extracting RNA using the TRIsure™ protocol (Bioline). Extractions from three replicates at each of the seven timepoints were used for a total of 21 RNA samples. RNA quality and quantity were qualitatively assessed using agarose gel electrophoresis. Samples were then DNase treated to remove any gDNA contamination (QuantSeq is sensitive to genomic DNA contamination) and RNA was re-extracted using the same TRIsure™ protocol. Quality and quantity were re-assessed using the Bioanalyzer 2100 (Agilent) before preparing libraries. Libraries were prepared using the QuantSeq 3’ mRNA-Seq Library Prep Kit FWD (Lexogen, Cat no. 015.24). Following library preparation, samples were analysed on a Bioanalyzer (Agilent 2100), Qubit fluorometric quantitation and qPCR to determine library concentration before sequencing. Samples were sequenced using single-end 75 bp chemistry on the NextSeq 500 (Illumina). Three additional sea anemone datasets (*Diadumene lineata, Stichodactyla mertensii* and *Triactis producta*) were also generated, to be used in comparative ortholog analysis (methods section: ‘Evolution of genes involved in regeneration’). Datasets were produced using the following methods: Total RNA was extracted from whole organisms by first homogenizing individuals in liquid nitrogen, followed by a TRIzol/chloroform RNA extraction protocol (TRIzol®, Life Technologies). Extracted RNA was assessed for quality and integrity on a Bioanalyzer 2100 (Agilent) using an RNA nano chip. Sequencing libraries were prepared in-house using the Illumina TruSeq® Stranded mRNA Library Preparation Kit and sequenced by Novogene using 150 bp paired-end chemistry on an Illumina HiSeq 4000. Transcriptomes were assembled using Trinity *de novo* assembler version 2.2.0 (Haas et al. 2013) using the Trimmomatic flag (Bolger et al. 2014) to trim and remove low quality reads. Cd-Hit version 4.6.4 was used to remove redundant sequences by clustering contigs with sequence similarity above 95%.

### Sequencing analysis, read trimming and alignment

Initial quality control results were provided by BaseSpace (Illumina) for all sequenced libraries and this showed the percentage of reads >Q30. Reads were trimmed and poly-A tails removed using a custom R pipeline that primarily uses the package ShortRead (https://rpubs.com/chapmandu2/171024). Samples were aligned to the *Exaiptasia* genome v1.1 using Bowtie2. As *E. pallida* undergoes symbiosis with *Symbiodinium* species, all reads that did not align, including potential contamination from *Symbiodinium* sp. were discarded. BAM files were then uploaded to Galaxy-QLD and a DGE count matrix was generated using HTSeq with default settings with the following changes: samples were specified as stranded and the transcript name was used as the ID setting. All trimmed reads were also aligned to the *Symbiodinium minutum* genome v1.0 (Shoguchi et al. 2013) and a count matrix produced using the same method as above.

### Differential gene expression (DGE) analysis

#### Count matrices and heatmaps

The count matrices (from *Exaiptasia* and *Symbiodinium*) produced by HTSeq was then used for differential count analysis using edgeR (Robinson et al. 2010). Pairwise sample differential expression analysis was performed using custom R scripts (https://github.com/zkstewart) at a p-value of 0.01, using TMM normalised triplicate samples. All of the following analyses were performed for *Exaiptasia* only. Pairwise comparisons were then annotated with GO terms, gene names and product names extracted from the available genome annotation. Pfam annotations were generated using a HMMer search against the Pfam database [version 31.0] with an E-value cut-off of 1 × *e*^−5^.. For further analysis of the differentially expressed gene set, another count matrix was generated containing only differentially expressed genes. Principle component analysis (PCA) plots, sample correlation matrix and heatmaps were generated from this DGE matrix using Perl-to-R scripts from Trinity (https://github.com/trinityrnaseq/trinityrnaseq/wiki/QC-Samples-and-Biological-Replicates). Parameters for each plot were: for heatmaps, --heatmap --log2 -- center_rows --gene_dist euclidean --sample_dist euclidean --heatmap_scale_limits “<2,2”. For PCA plots, --prin_comp --log2. For sample correlation matrix --sample_cor.

#### Subcluster analysis and GOseq

Subcluster analysis was performed on all differentially expressed genes (DEGs) using the Trinity perl script define_clusters_by_cutting_tree.pl and defined by cutting the tree based on 50 % max height (-Ptree 50). Subclusters show the centred log2 fold change of differentially expressed genes over the timecourse. Subclusters provide insight into transcriptional trends across the entire timecourse and provide a level of validation for the datasets, i.e., trends such as an overall increase in transcription factor activity can be expected to occur.

GO terms were extracted from the *Exaiptasia* genome v1.0 assembly and matched to the same gene IDs in v1.1. All genes that were differentially expressed across timepoints were split into separate upregulated and downregulated files. The Trinity perl script run_GOseq.pl was used to output depleted and enriched GO terms for each of the ten subclusters. An FDR cut-off of 0.05 was considered significant for depleted or enriched GO terms.

### Timecourse analysis

For further analysis of specific gene expression changes over timepoints, genes differentially expressed in the six pairwise comparisons between T0 vs. all other timepoints (T1-T6) were considered as the timecourse. Timepoints are referred to throughout this manuscript as “T1”, “T2” etc., rather than as “T0 vs T1”, “T0 vs T2” etc. To identify *Wnt* and *Wnt* pathway genes, all gene annotations were investigated and logFC for each gene was plotted. The list of *Wnt* pathway genes were compiled from those investigated by Schaffer *et al.* (2016), where they compared differential expression across axes in Planaria and *Nematostella.*

#### Immune system genes

To provide a more comprehensive understanding of the role of known innate immune genes in the regeneration response of *E. pallida* over the 72 hpa (hours post amputation), all genes annotated with the GO term ‘immune system process’ (GO ID: GO:0002376) were investigated.

#### Novel immune system genes

To analyse the role of novel innate immune genes in regeneration, the same method as used previously (van der Burg et al. 2016) was employed. TIR-containing genes were searched for using the Pfam IDs for TIR domain (PF01582), TIR_like domain (PF10137) and TIR-2 domain (PF13676). Cnidarian ficolin-like proteins (CniFl) were investigated by searching for genes with collagen Pfam domains (PF01391) in the annotations and searching accompanying Pfam annotations for any fibrinogen (PF00147) or Ig domains (Immunoglobulin superfamily CL0011 members includes: PF00047, PF07679, PF13927, PF13895, PF16681).

#### Evolution of genes involved in regeneration

In order to analyse the role of all novel and annotated *E. pallida* genes in regeneration, in the context of evolution, OrthoFinder (version 2.1.4) was used to find orthologous genes between the differentially expressed gene dataset (1547 unique genes) from *E. pallida* and the complete predicted peptidome from 27 datasets (from a total of 24 species, see Table 1 and Supplementary Table S7). All species FASTA files were prepared using two custom scripts (https://github.com/zkstewart) to prepare and run the files through BLAST. Orthologues were detected using the OrthoFinder python script (default settings) (Emms and Kelly 2015). Counts for the number of Orthogroups from each species in common with the *E. pallida* DGE dataset were obtained from CSV files (DGE_v_$species.csv) generated by OrthoFinder in output folder Orthologues_DGE. Species-specific novel genes in the *E. pallida* regeneration data set were identified from the ‘Orthogroups_UnassignedGenes.csv’ file.

**Table 1:**
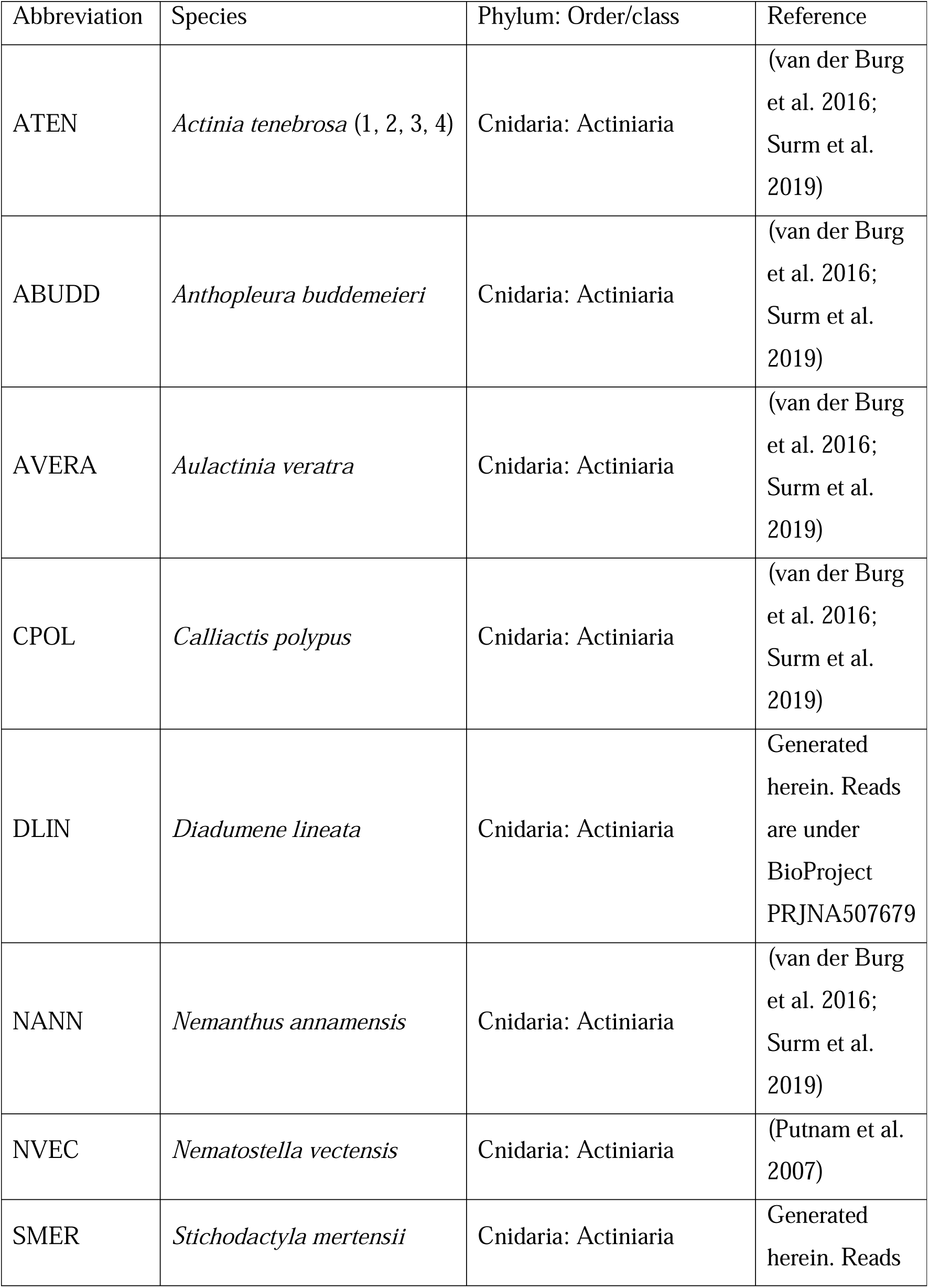
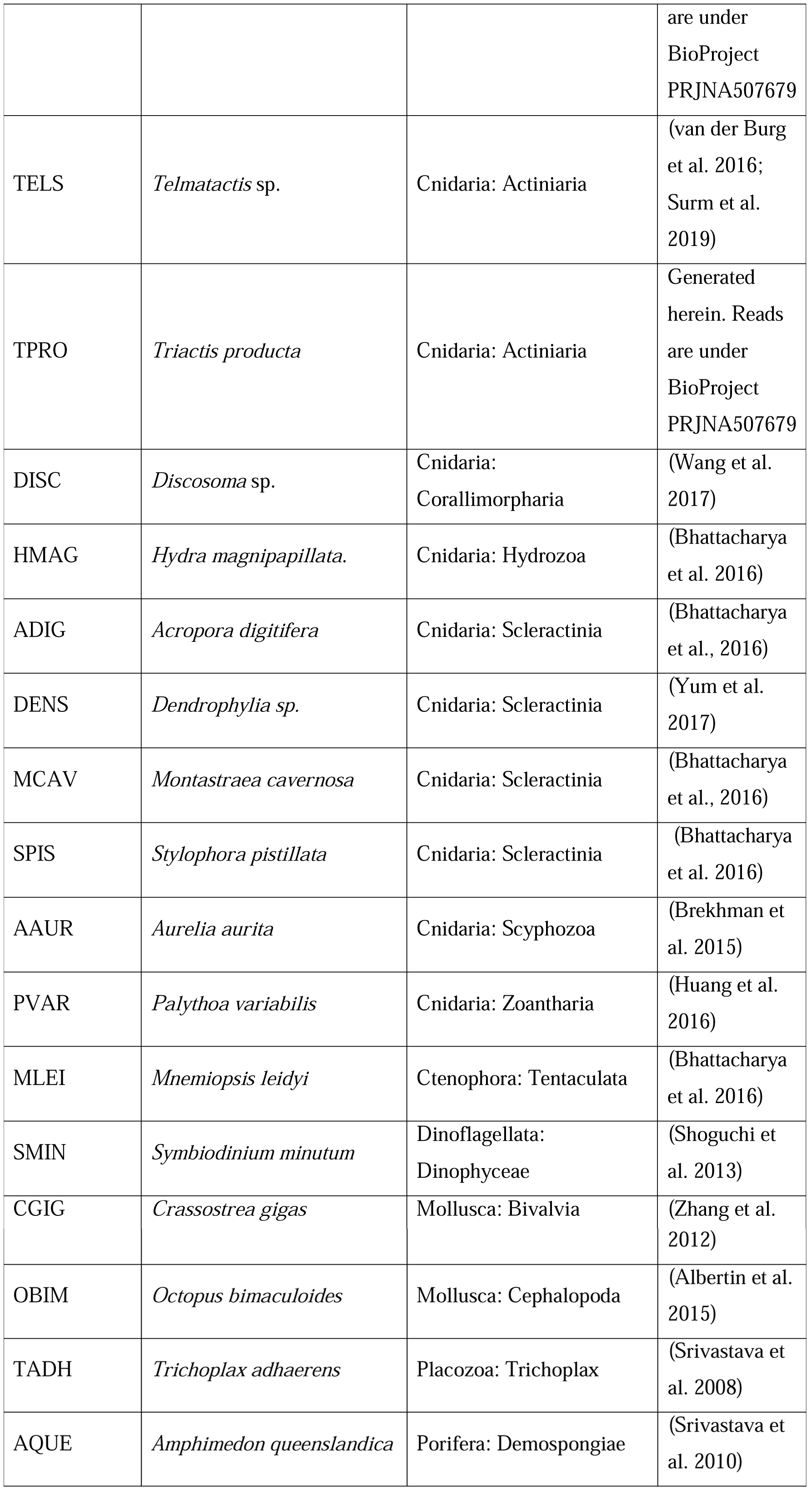
List of species used for OrthoFinder analysis. More information in Supplementary Table S7 describing the data source.

#### Validating species-specific novel genes

The 59 species-specific novel protein sequences were used as BLASTp queries against the predicted peptidomes of two *de novo* assembled *E. pallida* transcriptomes. Transcriptome #1 source: Sequence read archive SRX231866, run SRR696721, Aposymbiotic CC7 Transcriptome, assembled locally with Trinity. Transcriptome #2 source: Sequence read archive SRX757525: Aiptasia mRNA adult aposymbiotic, assembled locally with Trinity. BLASTp was run locally using blast+/2.6.0 with an E-value cut-off of 1e^−10^. The Trinity perl script analyze_blastPlus_topHit_coverage.pl was then used to obtain outputs showing the percentage coverage that each query sequence hit with the subject sequence. A conservative cut off of 85% was used for considering if a hit was valid, with the assumption that if genes with a very similar but not exact sequence match to the species-specific novel genes are being expressed in the selected transcriptomes then they are likely genes from the same family, and therefore the novel gene is likely a real gene within the genome. This cut-off also accounts for assembly errors in the two transcriptomes.

## Results

### Animal acquisition and visual observations

Multiple animals (n = 5) were filmed over several consecutive days, in order to observe tentacle and column regeneration. The main observations made of the regeneration process are: that *E. pallida* regenerates rapidly, i.e., within approximately 24 hours wound healing appears complete and the tentacle buds begin to emerge; that tentacle buds emerge from column striations at the site of amputation; that regeneration appears to be complete within 72-90 h (Supplementary File 1, Figures S1 and S2).

### Sequencing analysis, read trimming and alignment

The number of reads from each sequencing library ranged from 11,585,352 to 31,310,217 (x = 20,078,017 reads, σ = 4,665,526). QC results provided by BaseSpace (Illumina) for all sequenced libraries showed 85.4 % of all reads were >Q30. Read trimming reduced the number of reads per sample on average by 30.73% (σ = 0.0770), changes in read numbers for each dataset are shown in Supplementary Table S1a. Reads were aligned to the *Exaiptasia* draft genome scaffolds with an average success rate of 43.96% (σ = 0.0423). Reads were aligned to the *Symbiodinum* genome with an average success rate of 8.13% (σ = 0.0283). Three sea anemone transcriptomes were also generated and used for OrthoFinder analysis. For the three additional sea anemone datasets (*D. lineata. S. mertensii* and *T. producta*), assembly statistics can be viewed in Supplementary Table S1b.

### Differential gene expression (DGE) analysis

#### Count matrices and heatmaps

Of the 41,925 genes in the *Symbiodinum* genome, 30,195 (72.0%) genes were expressed at least once in all 21 datasets. No differential expression was detected for genes in the *Symbiodinium* genome (data not shown but the count matrix can be provided upon request). Of the 26,042 genes in the *Exaiptasia* genome, 25,209 (96.8%) genes were expressed at least once in all 21 datasets (Supplementary Table S2). Overall, 4,435 instances of differential expression occurred and many of these instances were the same gene being differentially expressed multiple times across pairwise comparisons. 1,547 unique genes were found to be significantly differentially expressed across all 21 pairwise comparisons. A count matrix with the 1,547 DEGs was generated to further analyse the dataset in greater detail (Supplementary Table S3). Principal component analysis (PCA), sample correlation and sample clustering of differentially expressed genes (Supplementary File 2, Figures S3-S5) showed that most samples cluster either with their designated timepoint or with the subsequent timepoint, indicating most samples are consistent replicates. DEGs were used to understand major transcriptional changes over the regeneration time course.

#### Subcluster analysis and GOseq

Ten subclusters were recovered from the heatmap of significantly differentially expressed genes, which each are comprised of a number of genes that all show similar expression patterns over the seven timepoints (Supplementary File 2, Figure S6-S7). Two of the subclusters (#4 – 469 transcripts and #7 – 205 transcripts) were found to have significantly enriched GO terms. In subcluster 4, most transcripts show an expression level maintained at a baseline level throughout the timecourse, with a spike in transcriptional activity at 2.5 hpa (T2). These transcripts were annotated with terms associated with cell communication, cell signalling and response to signal and stimulus, as well as regulation of metabolic, biological and cellular activity or processes (Table 3). In subcluster 7, transcript expression increases from a baseline level with a sudden drop in expression at 45 min – 2.5 hpa, followed by an increase in expression which generally maintains at a baseline level throughout the rest of the timecourse. These transcripts were annotated with terms associated with cytoskeletal remodelling, cellular movement and nucleic acid/base metabolic activity (Table 2). One subcluster (#4 – 469 transcripts) was found to have significantly depleted GO terms, which were terms associated with DNA replication and repair and nucleic acid metabolic activity (Table 4). The full list of GO terms (with GO IDs and FDR values) are in Supplementary Table S4. To visualise GO terms in subclusters 4 and 7, broad descriptions are displayed next to each subcluster graph (Figure 2).

**Table 2.** Subcluster 7 significantly <u>enriched</u> GO terms classified into broader group descriptions. BP = biological process, CC = cellular component, MF = molecular function. Group descriptions are not GO terms.

| GO term | Group description |
| --- | --- |
| CC microtubule | Cell movement and cellular remodelling |
| MF motor activity |  |
| CC cytoskeleton |  |
| CC cytoplasmic dynein complex |  |
| CC microtubule associated complex |  |
| CC axoneme part |  |
| CC cytoskeletal part |  |
| CC cilium |  |
| CC dynein complex |  |
| MF microtubule motor activity |  |
| CC ciliary part |  |
| CC axonemal dynein complex |  |
| BP microtubule-based process |  |
| BP microtubule-based movement |  |
| BP purine nucleoside triphosphate metabolic process | Energetics and nucleic acid building block metabolism |
| BP nucleoside triphosphate metabolic process. |  |
| BP nucleotide catabolic process |  |
| BP nucleoside phosphate catabolic process |  |
| BP organophosphate catabolic process |  |
| BP purine nucleotide catabolic process |  |
| BP purine-containing compound catabolic process |  |

**Table 3.** Subcluster 4 significantly <u>enriched</u> GO terms classified into broader group descriptions. BP = biological process, MF = molecular function. Group descriptions are not GO terms.

| GO term | Group description |
| --- | --- |
| BP regulation of small GTPase mediated signal transduction | Cell communication and cell signalling |
| BP regulation of signalling |  |
| BP neuropeptide signaling pathway |  |
| MF signal transducer activity |  |
| BP signal transduction |  |
| BP signal transduction by protein phosphorylation |  |
| BP regulation of cell communication |  |
| BP cell surface receptor signaling pathway |  |
| MF molecular transducer activity |  |
| BP regulation of signal transduction |  |
| BP transmembrane receptor protein tyrosine kinase signaling pathway |  |
| BP intracellular signal transduction |  |
| BP regulation of intracellular signal transduction |  |
| BP G-protein coupled receptor signaling pathway |  |
| MF receptor activity |  |
| BP small GTPase mediated signal transduction |  |
| MF signaling receptor activity |  |
| MF transmembrane signaling receptor activity |  |
| BP cellular response to chemical stimulus | Response to stimulus or substance |
| BP response to stimulus |  |
| BP cellular response to stimulus |  |
| BP positive regulation of response to stimulus |  |
| BP regulation of response to stimulus |  |
| BP cellular response to growth factor stimulus |  |
| BP response to endogenous stimulus |  |
| BP cellular response to organic substance |  |
| BP response to growth factor |  |
| BP response to organic substance |  |
| BP response to chemical |  |
| BP cell differentiation | Cell proliferation or differentiation |
| BP positive regulation of stem cell proliferation |  |
| BP regulation of phosphorus metabolic process | Regulation of phosphate activity |
| BP regulation of phosphate metabolic process |  |
| BP positive regulation of phospholipase C activity |  |
| BP regulation of phospholipase C activity |  |
| BP MAPK cascade | MAPK cascade |
| BP regulation of MAPK cascade |  |
| BP lung-associated mesenchyme development | Lung/gland changes |
| BP lung development |  |
| BP gland morphogenesis |  |
| BP biological regulation | Regulation of activity or processes |
| BP regulation of catalytic activity |  |
| BP regulation of biological process | (biological/cellular/metabolic) |
| BP regulation of metabolic process |  |
| BP regulation of cellular process |  |
| BP positive regulation of cellular process |  |
| BP regulation of molecular function |  |
| BP regulation of cellular metabolic process |  |
| BP regulation of primary metabolic process |  |
| BP regulation of hydrolase activity |  |

**Table 4.** Subcluster 4 significantly <u>depleted</u> GO terms classified into broader group descriptions. BP = biological process, MF = molecular function. Group descriptions are not GO terms.

| GO term | Group description |
| --- | --- |
| CC microtubule organizing center | DNA replication |
| BP DNA replication |  |
| MF DNA polymerase activity |  |
| MF nucleotidyltransferase activity | DNA repair |
| BP RNA-dependent DNA biosynthetic process |  |
| MF RNA-directed DNA polymerase activity |  |
| MF RNA binding | General nucleic acid metabolic process |
| MF catalytic activity |  |
| MF nucleic acid binding |  |
| BP DNA metabolic process |  |

**Fig. 2.**
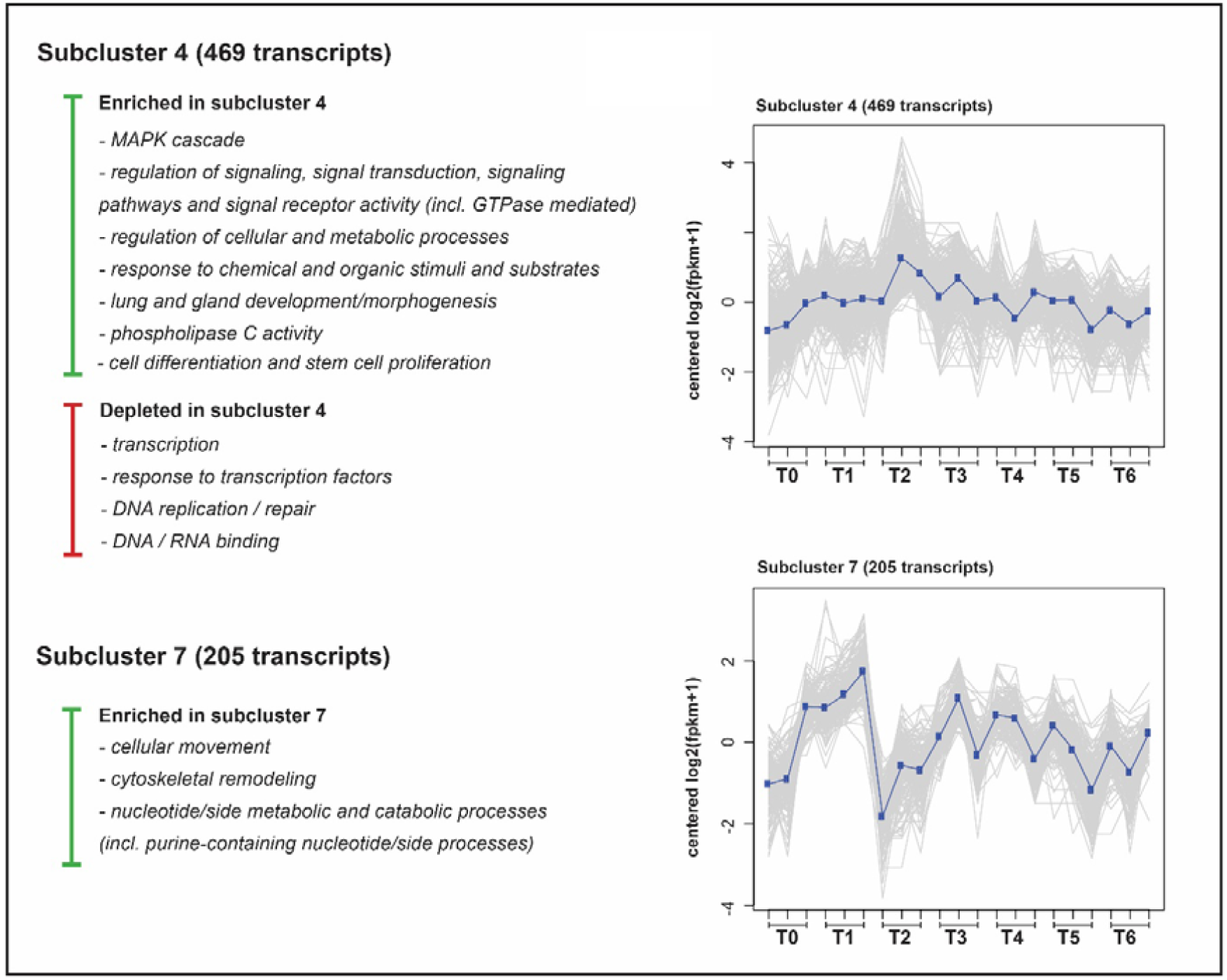
Subclusters determined to have significantly enriched and depleted GO terms as determined by GOseq to an FDR of 0.05. Ten subclusters were recovered from the heatmap of differentially expressed genes, but only two were found to have genes annotated with GO terms that are significantly enriched or depleted. Broad classifications and all significant GO terms can also be viewed in tables 2, 3 and 4.

### Timecourse analysis

A further fine-scale analysis of specific gene expression changes over timepoints was performed. Only the genes differentially expressed in the six pairwise comparisons between the baseline T0 to all other timepoints (T1-T6) were analysed, henceforth referred to as the timecourse. A total of 932 instances of differential expression occurred across the timecourse, consisting of 625 unique genes (several genes were DE more than once). Figure 3 shows a summary of the number of DEGs and their annotations. Figure 4 shows the log fold change (logFC) of each of the 932 significantly differentially expressed genes, arranged in order of highest to lowest logFC within each timepoint. The seven *Wnt* and *Wnt* pathway related genes that are expressed during the timecourse are shown on this figure, which are all upregulated across T2 to T5.

**Fig. 3.**
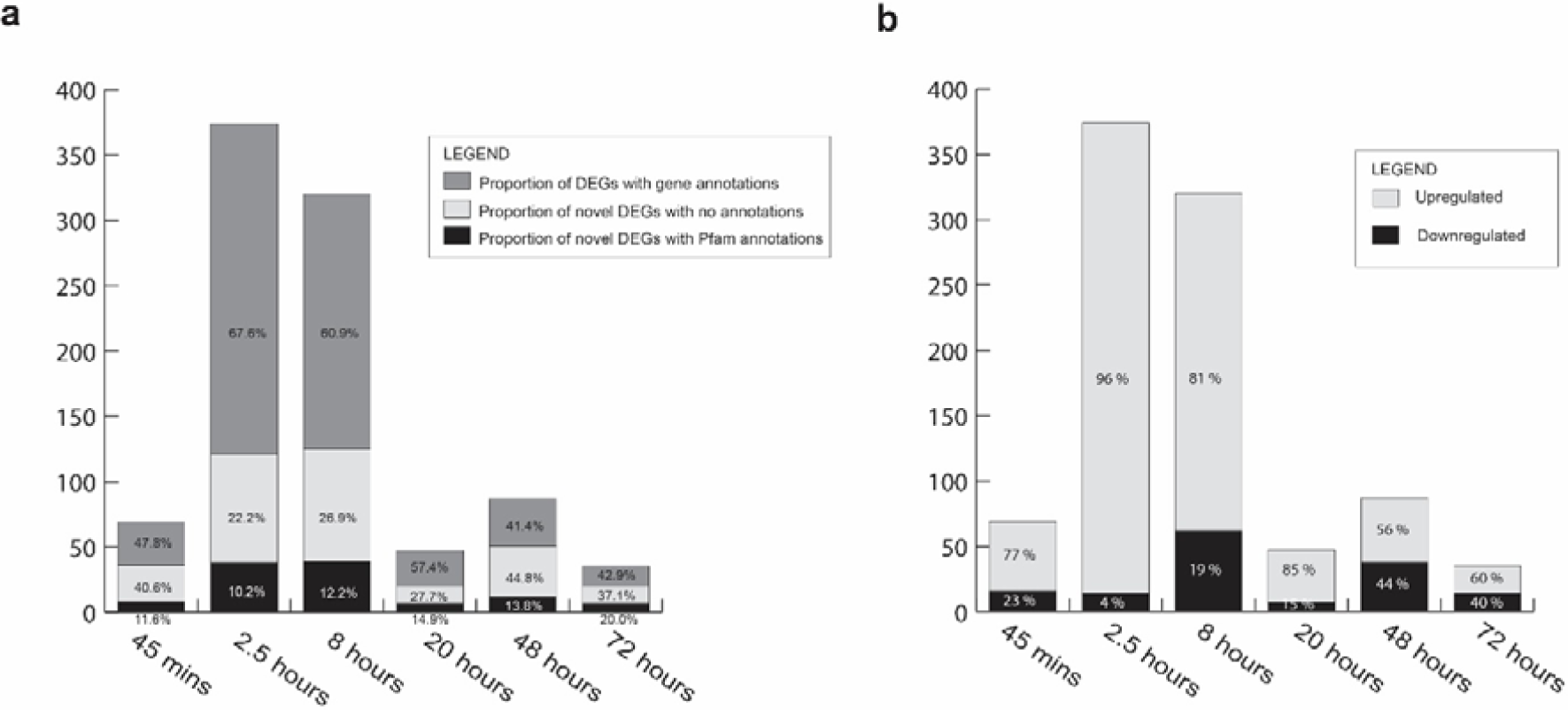
Proportions of differentially expressed genes (DEGs) with/without annotations and up or down regulated. Graph **a)** shows the number of DEGs at each time point and the proportion of these genes that have gene annotations (known gene names based off BLAST hits), the proportion of novel genes (i.e., no annotations at all) and the proportion of novel genes with at least one Pfam annotation. T1 (45 mpa) = 69 DEGs; T2 (2.5 hpa) = 374 DEGs; T3 (8 hpa) = 320 DEGs; T4 (20 hpa) = 47 DEGs; T5 (48 hpa) = 87 DEGs; T6 (72 hpa) = 35 DEGs. Graph **b)** shows the number of DEGs at each time point and the proportion of these genes that are up- or down-regulated.

**Fig. 4.**
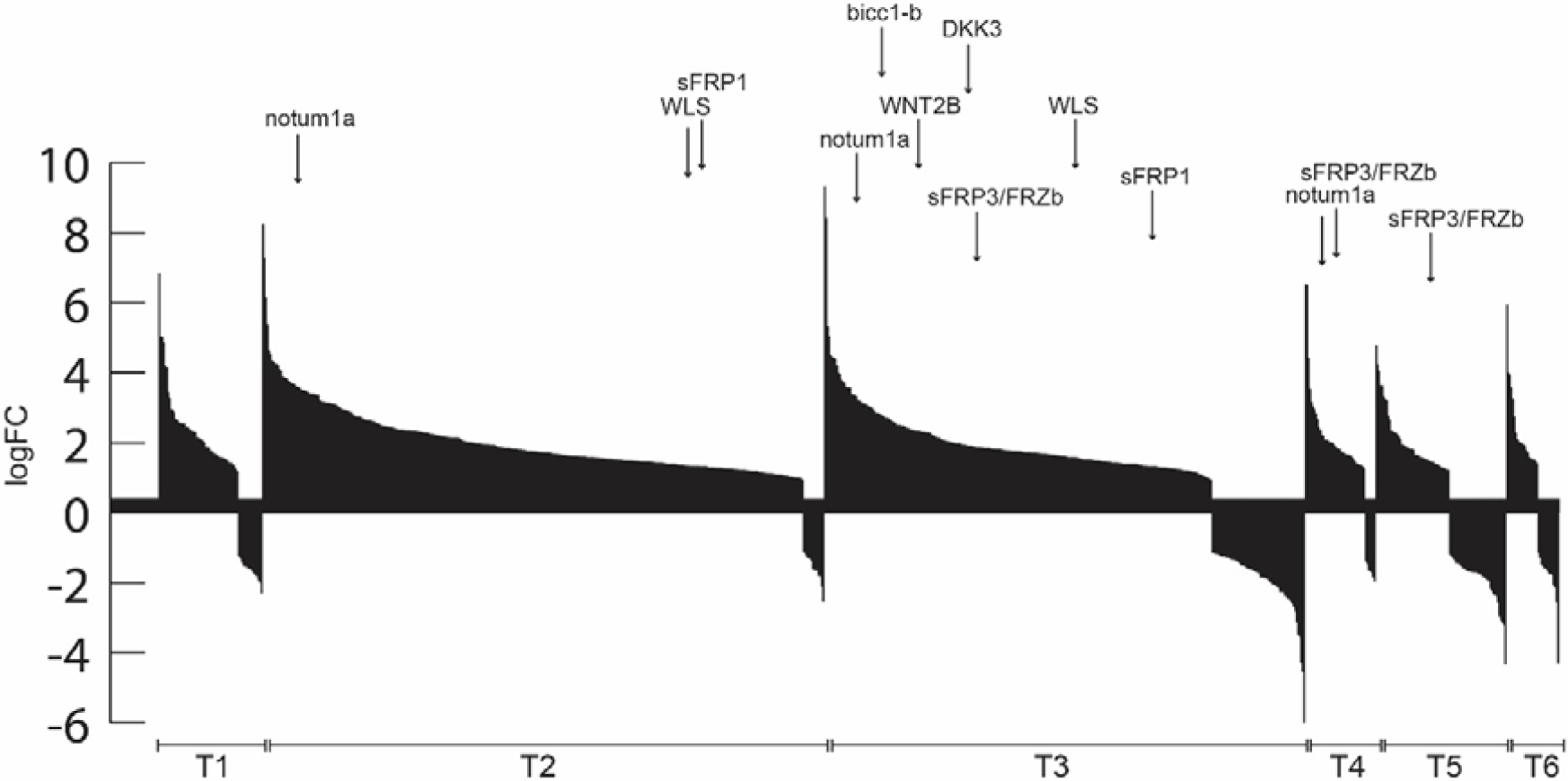
Column graph depicts the log fold change (logFC) of each significantly differentially expressed gene at each timepoint. The 932 genes are arranged within each timepoint in order from highest logFC to lowest logFC. **T2**: gene *WLS* (gene10121) logFC = 1.26; *notum1a* (gene22172) logFC = 3.75; *sFRP1* (gene8137) logFC = 1.23. **T3**: gene *WNT2B* (gene6631) logFC = 2.50; *WLS* (gene10121) logFC = 1.51; *notum1a* (gene22172) logFC = 3.36; *bicc1-b* (gene26408) logFC = 2.78; *DKK3* (gene9249) logFC = 2.04; *sFRP3/FRZb* (gene19466) logFC = 1.98; *sFRP1* (gene8137) logFC = 1.31. **T4**: *notum1a* (gene22172) logFC = 2.66; *sFRP3/frzb* (gene19466) logFC = 1.71. **T5**: *sFRP3/FRZb* (gene19466) logFC = 1.37. No *WNT* or *WNT* pathway associated genes were downregulated. None were found in T1 or T6.

#### Immune system genes

To examine the expression of the immune system genes in the regeneration process, the “immune system process” GO term (GO:0002376) was used to find differentially expressed innate immune genes. These are listed below and can also be found in Supplementary Table S6 which shows the full annotation data for these immune genes. To understand the immune response in context with other major processes, the number of genes associated with multiple major processes are represented alongside all immune gene counts in Figure 5a and 5b. At 45 mpa (minutes post amputation) (T1), two differentially expressed genes (ETS domain-containing protein Elk-1 (+) and TNF receptor-associated factor 3 (+) are annotated with the immune system process GO term, both are upregulated (+). This represents 2/69 (2.90 %) of the differential expression at this timepoint.

**Fig. 5.**
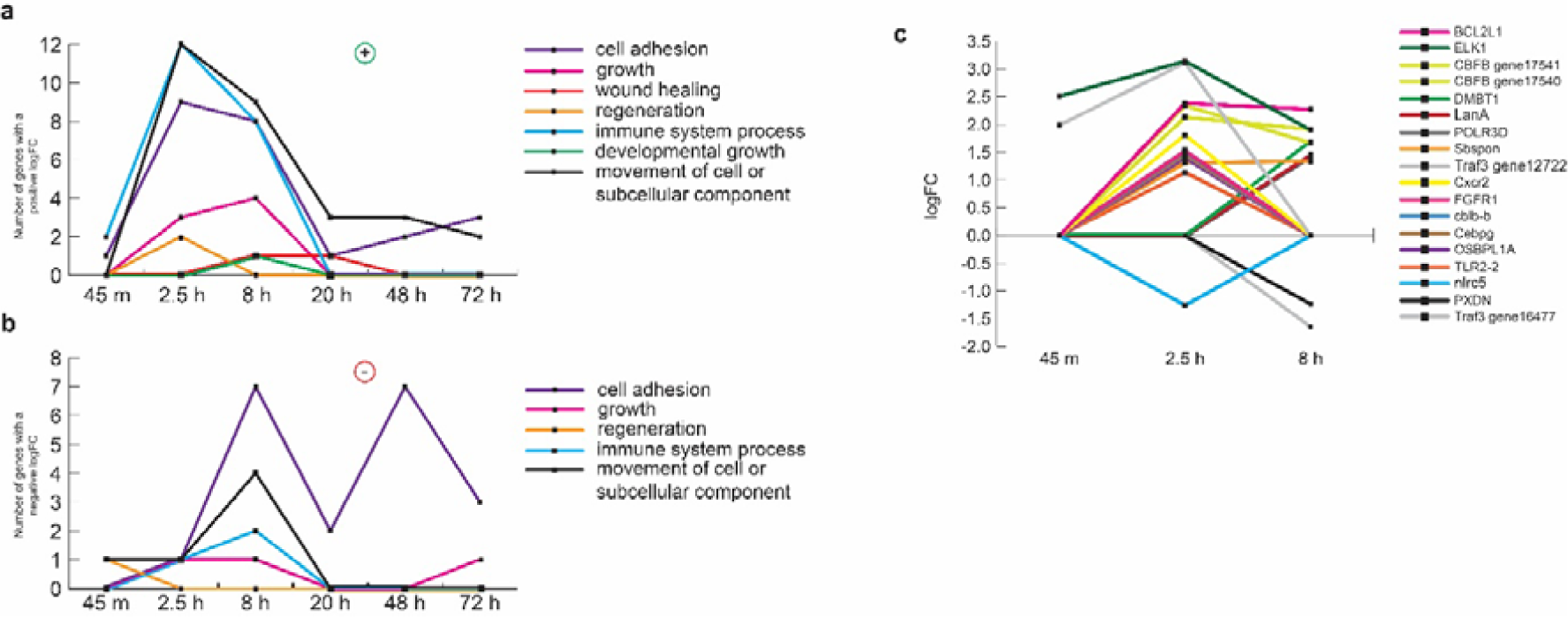
The immune system process in context. Graphs **a)** and **b)** depict the number of up- and down-regulated genes (respectively) annotated with GO terms in major categories at each timepoint. GO terms are as follows: ‘Cell adhesion’: GO:0007155; ‘Movement of cell or subcellular component’: GO:0006928; ‘Developmental growth’: GO:0048589; ‘Wound healing’: GO:0042060; ‘Immune system process’: GO:0002376; ‘Regeneration’: GO:0031099; ‘Growth’: GO:0040007. **c)** Genes annotated with the immune system process GO term. Timepoints 4, 5 and 6 are not shown as no genes were annotated with the GO term past 8 hours (T3).

At 2.5 hpa (T2), there are 13 differentially expressed immune genes, with 12 upregulated (+) and one downregulated (-). These are: ETS domain-containing protein Elk-1 (+), TNF receptor-associated factor 3 (+), Bcl-2-like protein 1 (+), core-binding factor subunit beta, two copies (+), C-X-C chemokine receptor type 2 (+), Fibroblast growth factor receptor 1 (+), E3 Ubiquitin-protein ligase CBL-B-B (+), CCAAT/enhancer-binding protein gamma (+), Oxysterol-binding protein-related protein 1 (+), Somatomedin-B and thrombospondin type-1 domain-containing protein (+), Toll-like receptor 2 type-2 (+), and protein NLRC5 (-) This represents 13/374 (3.48 %) of the differential expression at this timepoint.

At 8 hpa (T3), there are 10 significantly differentially expressed genes, with eight upregulated (+) and two downregulated (-). These are: Bcl-2-like protein 1 (+), core-binding factor subunit beta (+), ETS domain-containing protein Elk-1 (+), Deleted in malignant brain tumors 1 protein (+), core-binding factor subunit beta (+), Laminin subunit Alpha (+), DNA-directed RNA polymerase III subunit RPC4 (+), Somatomedin-B and thrombospondin type-1 domain-containing protein (+), Peroxidasin-like (-) and TNF receptor-associated factor 3 (-). This represents 10/320 (3.13 %) of the differential expression at this timepoint.

These immune genes are represented in Figure 5c. From 20 hours on to 72 hours (i.e., T4, T5 and T6) no genes are annotated with the GO term for immune system process.

#### Novel immune system genes

We analysed the possible role of novel immune genes in regeneration for all differentially expressed genes that were annotated as hypothetical proteins (as predicted from the genome assembly). Using a previously described method, TIR domains (Toll/interleukin-1 receptor homology domain) was used to find potential novel immune genes.

No TIR domain (PF01582) Pfam annotations were found in any of the differentially expressed hypothetical proteins. One annotated gene ‘Toll-like receptor 2 type-2’ (gene1475) was found containing the TIR domain (PF10582) and is upregulated at 2.5 hpa (T2). However, this gene name annotation is likely incorrect as the protein does not have the typical Pfam domains of a TLR (canonical TLR contain multiple LRR, a TMD and a TIR domain) but rather has the typical Pfam domains of an Interleukin-like receptor (multiple Ig domains, a TMD and a TIR domain).

One hypothetical gene (gene25976 | KXJ21747.1 | AIPGENE15161) was found containing the TIR-2 domain (PF13676) and an Armadillo (ARM)/beta-catenin-like repeat, which was found just upstream of the TIR-2 domain but at a lower E-value (1e-4) threshold and was up-regulated at 2.5 hpa (T2). At 8 hpa (T3) and at 48 hpa (T5) the gene is downregulated.

No TIR_like domains (PF10137) were found in any hypothetical or annotated differentially expressed genes.

Two potential Cnidarian ficolin-like genes (*CniFl*) were identified, both are differentially expressed at 8 hpa (T3), one is upregulated and one is downregulated. Both are annotated as Ryncolin genes, which typically have collagen and fibrinogen domains but not Ig domains, which are hallmarks of CniFls. Both potential CniFl genes have one collagen, multiple Igs, and a fibrinogen domain. Gene12803 (KXJ26376.1 | AIPGENE11648) and gene11981 (KXJ13472.1 | AIPGENE13580) both have identical Pfams annotations; collagen (PF01391), 4 Igs (PF13927, PF07679, PF13895, PF00047), and fibrinogen_C (PF00147).

#### Evolution of genes involved in regeneration

OrthoFinder assigned 1,313,495 genes (84.5% of total) to 132,195 orthogroups. Fifty percent of all genes were in orthogroups with 27 or more genes (G50 was 27) and were contained in the largest 9907 orthogroups (O50 was 9907). There were 34 orthogroups with all species present and none of these consisted entirely of single-copy genes.

The counts of the number of orthogroups from each species in common with the differentially expressed dataset from *E. pallida* were recorded and plotted in Figure 6. This figure shows an increasing trend of more orthogroups shared between the differential expression dataset and species more closely related to *E. pallida*. There is a large increase in the number of shared orthogroups between the Ctenophore *Mneiopsis leidyi* (∼100 shared orthogroups) and the sponge *Amphimedon queenslandica* (∼500). This trend increases upwards through Metazoan and Cnidarian species, with a noticeable drop at *H. magnipapillata*. The trendline is observed to plateau through the anemone species at a high number of shared orthogroups (∼950), although the model anemone species *N. vectensis* is observed to have lower shared orthogroups (∼ 800) than all other anemones.

**Fig. 6.**
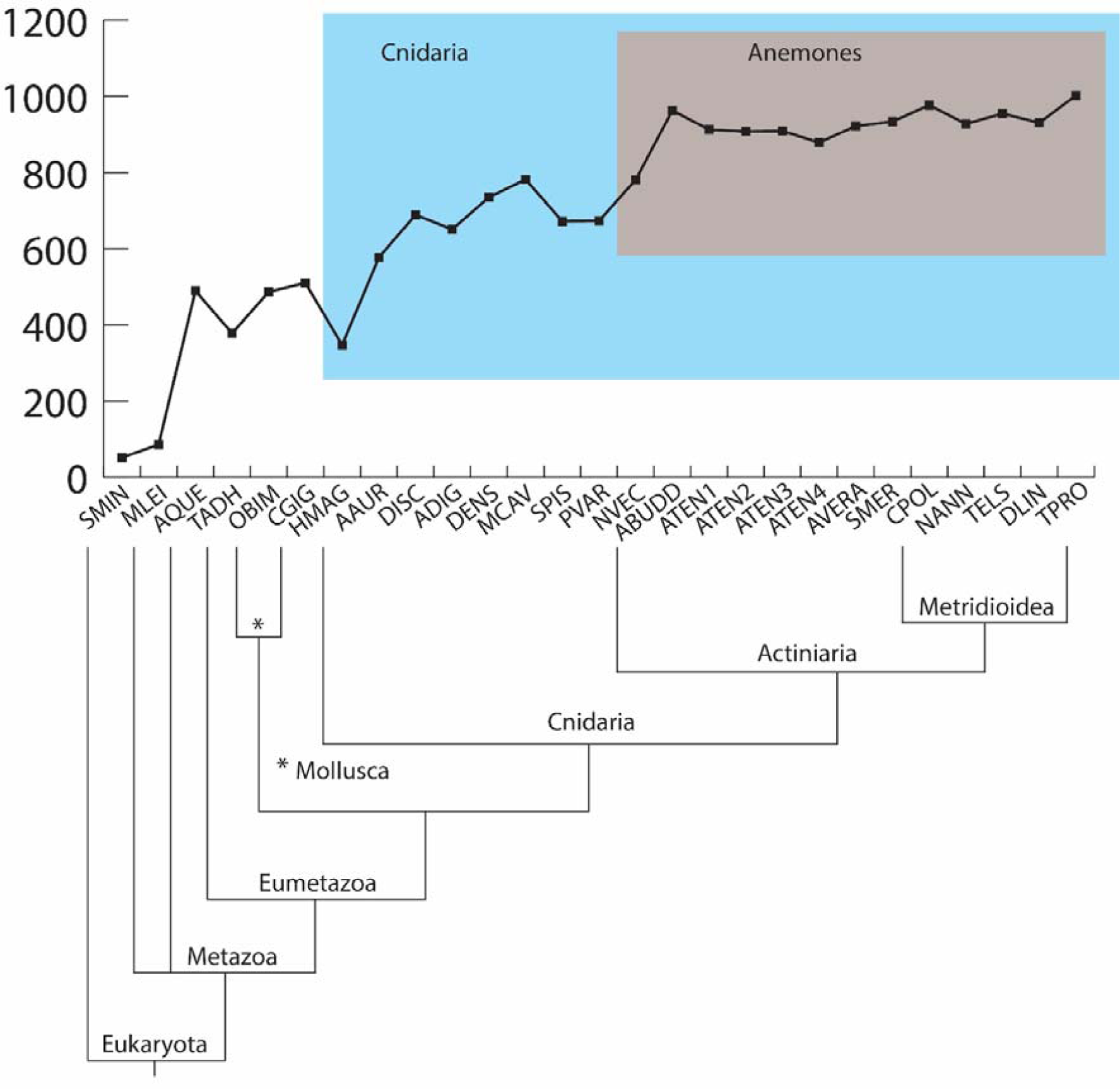
Number of orthogroups shared between the gene set of each species compared with the differentially expressed gene set from *Exaiptasia pallida*. Species are arranged in order of phylogenetically most distant from *E. pallida* (leftmost species on the x-axis) to phylogenetically most close to *E. pallida* (rightmost species on the x-axis). Codes for species names are as follows: SMIN *Symbiodinium minutum*; MLEI *Mnemiopsis leidyi*; AQUE *Amphimedon queenslandica*; TADH *Trichoplax adhaerens*; OBIM *Octopus bimaculoides*; CGIG *Crassostrea gigas*; HMAG *Hydra magnipapillata*; AAUR *Aurelia aurita*; DISC *Discosoma* sp.; ADIG *Acropora digitifera*; DENS *Dendrophylia* sp.; MCAV *Montastraea cavernosa*; SPIS *Stylophora pistillata*; PVAR *Palythoa variabilis*; NVEC *Nematostella vectensis*; ABUDD *Anthopleura buddemeieri*; ATEN1 *Actinia tenebrosa* (red); ATEN2 *Actinia tenebrosa* (brown); ATEN3 *Actinia tenebrosa* (green); ATEN4 *Actinia tenebrosa* (blue); AVERA *Aulactinia veratra*; SMER *Stichodactyla mertensii*; CPOL *Calliactis polypus*; NANN *Nemanthus annamensis*; TELS *Telmatactis* sp.; DLIN *Diadumene lineata*; TPRO *Triactis producta*

#### Species-specific novel genes

Fifty-nine of the differentially expressed genes in *E. pallida* are not orthologues to any gene from the other investigated species and are henceforth referred to as the 59 species-specific novels (listed in Supplementary Table S8a). Only 23 species-specific novels obtained BLAST hits to ORFs in either of the *E. pallida* transcriptomes, indicating the remaining 36 are either not expressed in these transcriptomes or are products of incorrect gene model annotation. Of these 23, only 10 species-specific novels had a hit that covered at least 85% of the length of the subject. Therefore, these 10 are the most confidently predicted to be true species-specific genes in *E. pallida*. Nine are annotated as hypothetical genes and one is annotated as a Collagen alpha-2(I) chain protein (KXJ17799.1 | AIPGENE23815 | gene5286). The other proteins in the 59 species-specific are all hypothetical proteins, except for one Kinesin-like protein KIFC3 (KXJ16016.1 | AIPGENE25729 | gene8131) and one other Collagen alpha-1(I) chain protein (KXJ17778.1). See Supplementary Tables S8a-d for full list of genes and BLAST results. Table S8e shows that the most confidently predicted novel *E. pallida* genes are expressed throughout the regenerative timecourse.

## Discussion

This present study is the first transcriptional description of regeneration in *Exaiptasia pallida*, a recently emerged model sea anemone species (Baumgarten et al. 2015). The aim was to understand the rapid regeneration response of *E. pallida* with a focus on the role of innate immune genes in this process. How gene expression in *E. pallida* compares to other evolutionary close relatives and regeneratively competent species was also of key interest.

The regenerative capacity of animals can vary significantly even among closely related species (Fumagalli et al. 2018; Liu et al. 2013; Tiozzo and Copley, 2015). Sea anemones are no exception to this and differences in regenerative strategy have been observed here and in other studies. *Nematostella vectensis* despite being the dominant model sea anemone species is highly divergent and sits in the small outgroup family Edwardsiidae, suborder Anenthemonae (order: Actiniaria) which is a separate suborder from most other anemones residing in the suborder Enthemonae (order: Actiniaria) (Putnam et al. 2007; Rodríguez et al. 2014). This difference is exemplified in this study in Figure 6; the anemone species with the least number of orthologous genes to the *E. pallida* DGE dataset is *N. vectensis.* This indicates that *N. vectensis* is not the most representative model species for anemone regeneration. Multiple recent studies have explored regeneration in *N. vectensis* from a transcriptional, morphological and cellular perspective (Amiel et al. 2015; DuBuc et al. 2014; Schaffer et al. 2016) and some similarities and differences can be drawn between this species and *E. pallida.*

Key genes found by Schaffer *et al.* (2016) to be differentially expressed in orally regenerating *N. vectensis* include multiple components of the *Wnt* pathway, multiple transcription factors and homeobox genes, consistent with results here for *E. pallida* and with an earlier study on *N. vectensis* (DuBuc et al. 2014). The homeobox gene *Otxc* was not found in the present study, consistent with it being found to denote physa (foot) not oral regeneration in *N. vectensis*. Many types of metalloproteinases (MP, including Zinc MP and *ADAMTS*) were found more highly expressed in orally (vs. aborally) regenerating *N. vectensis*, which is also seen in the present study across all timepoints (see Supplementary Tables S5), although whether these are more highly expressed in *E. pallida* in the oral side versus the aboral side is unknown. The paired-type homeobox gene *Dmbx*, which is upregulated in oral *N. vectensis* regeneration but downregulated in *Hydra* head regeneration (Petersen et al. 2015), is not present in *E. pallida. Hydra* and *N. vectensis* both show late oral expression of *Otp* and *Sox14*; *Otp* is not differentially expressed in *E. pallida* but *Sox14* is expressed (but only significantly when comparing T1 vs T3, Supplementary Table S5g). Few components of the BMP pathway were found, except for a *Tolloid-like protein* (astacin-like zinc-dependent metalloprotease) found at T3, and a *Noggin* gene at T3 and T4, which is fairly consistent with *N. vectensis*. In contrast to *N. vectensis, Notch* pathway genes were not differentially expressed in this study (except for one *notch1* gene DE at T1 vs T5, Supplementary Table S5g). There is no evidence that any of the DE *E. pallida* genes have either a chitin binding or synthesising protein domain (chitin genes were found highly enriched in *N. vectensis* (Schaffer et al. 2016)), although the GO term associated with ‘chitin binding’ (GO:0008061) is found in several genes with carbohydrate binding domains, one of which is Chitotriosidase-1 (DE at T1 vs T2, Supplementary Table S5g). Other results for *N. vectensis* found by DuBuc *et al.* (2014) are also consistently found in *E. pallida*, e.g., the MAPK cascade was proposed to be essential for would healing and is significantly enriched here and many metalloproteinases can be found throughout the timecourse. However, not all key results found by DuBuc et al. (2014) are found to be expressed in *E. pallida*, including *SoxE1* (several other *Sox* genes are expressed in *E. pallida* though), thiamine enzyme and maltase enzyme.

A single early transcriptional activity burst, comprised of genes involved primarily in cellular movement, remodelling, and communication characterise the response in both *E. pallida* and the phylogenetically close relative *Calliactis polypus* (Enthemonae: Metridioidea). A recent study explored the transcriptional response of *C. polypus* during a regenerative timecourse (Stewart et al. 2017). While the dissection method for *C. polypus* was different to this study (vertical vs transverse), *C. polypus* is observed to follow a very similar transcriptional trajectory to *E. pallida*, in terms of the observed initial early burst of activity, which then gradually drops and returns to baseline expression levels within a few days. The difference is observed at the specific time scale, as the major burst of activity is at 20 hpa, as opposed to 2.5 hpa in this study. Despite the small difference in time scale, the key findings of Stewart *et al.* (2017) are very similar to what we also find in *E. pallida*. While *Wnt* genes appear to be a feature of almost all animal regeneration studies (Chera et al. 2009; Duffy et al. 2010; Fumagalli et al. 2018; Schaffer et al. 2016), the lack of *Wnt* genes in the *C. polypus* study is likely due to the vertical dissection method, as *Wnt* expression generally denotes axis polarity in an organism (Manuel 2009). Additionally, a pulsing like motion was observed in *C. polypus*, which is also seen in *E. pallida* (Video file 1 and Supplementary File 1, Figure S2). This may be a similar mechanism as described in a recent paper on the symmetrisation of amputated jellyfish ephyra (Cnidaria: Scyphozoa: *Aurelia aurita*), which use a mechanical pulsing movement to regain symmetry in ephyra in order to regenerate (Abrams et al. 2015).

*Exaiptasia pallida* exhibits some aspects of morphallactic regeneration based on the initial visual observations here (Supplementary File 1, Figure S1), which show the individuals oral disc and tentacles regenerating from the amputation location, at least in the early stages of regeneration. The regeneration mechanism in *Hydra* is traditionally described as morphallactic, that is, regeneration can proceed through the remodelling of existing tissue only and with no input from cell proliferation (Morgan 1901; Bosch 2007), although some studies propose that regeneration in *Hydra* is highly variable and some regenerative strategies do rely cell proliferation (Miljkovic-Licina et al. 2007; Chera et al. 2009; Buzgariu et al. 2018). It is likely that after the initial wound closure and regeneration of the oral components that the animal essentially reverts back to a growth-like state. While it is yet to be confirmed if *E. pallida* indeed can regenerate without cell proliferation i.e., morphallactically, data presented here show few enriched GO terms associated with cell proliferation, suggesting it may not be a major contributor to the regeneration response. There is also not a large response of genes associated with ‘growth’ GO term in the later timepoints, so perhaps either this occurs later than 72 hpa, or the transcripts required to signal growth are expressed early in regeneration and this drives normal growth later on. Notably, cell proliferation and apoptosis are also not a feature of ephyra regeneration (Abrams et al. 2015), but perhaps this is due to either the age of the organism (i.e., the ephyra is the juvenile jellyfish) or due to the small size which allows ephyra to regenerate through symmetrization alone.

Many components of regeneration in other species are conserved and reflected in *E. pallida.* Fumagalli *et al.* (2018) performed a meta-analysis on *Hydra magnipapillata, Schmidtea mediterranea* (Platyhelminthes: Planaria), and *Apostichopus japonicus* (Echinodermata, sea cucumber) as well as some additional limb and liver regeneration datasets from axolotl and mammalian models, respectively, to understand the differences and similarities in regenerative strategies across taxa (Fumagalli et al. 2018). Fumagalli et al. (2018) highlighted the differences in time scales for regeneration time in different animals but managed to reconcile this by splitting the gene expression profiles into transient or sustained up- or down-regulation. The key outcomes of the Fumagalli et al (2018) study show in the early phases that upregulated GO terms were ‘cell-cell communication’, ‘DNA repair’, and ‘initiation of transcription’. The downregulated GO terms were ‘metabolic processes’, ‘cell adhesion’ and ‘immune-response-related genes’ (including cell adhesion, receptor mediated endocytosis, cellular component biogenesis, lipid metabolic process). Some results are noticeably different to our results. DNA repair is a depleted GO term (Figure 2 and Table 4), suggesting that DNA repair, along with other DNA metabolic activities, is absent in *E. pallida* regeneration, in contrast to perhaps many other species. Cell adhesion is a term that is noticeably absent from any significantly enriched or depleted activities, as detected during subcluster analyses. However, a general search for cell adhesion shows that it does appear throughout the timecourse and follows a similar trajectory to GO term for ‘movement of cell or subcellular component’ although after T3 ‘cell adhesion’ is largely represented in downregulated terms (Figure 5a and 5b).

The immune system in *E. pallida* plays only an early transcriptional role in the regeneration response. In terms of the number of differentially expressed genes, immune genes contribute a small percentage (2.9-3.5 %) to the differential expression at each timepoint. Figure 5 shows that in context, the immune system appears to play a comparably substantial role as other major processes ‘Movement of cell or subcellular component’ and ‘Cell adhesion’ in terms of the number of genes expressed that are annotated with these GO terms. However, after 8 hpa immune genes are not present at all. Immune genes are primarily upregulated, with only a couple downregulated in the early timepoints. During wound healing there is some requirement for the immune system to protect against infection, which likely explains the initial response in the first few hours. It should also be noted that while the ancestral GO term “immune system process” was used to classify immune genes, some genes detected in the current study, such as Bcl-2 and Elk-1, are known to have many other functions e.g. Bcl-2 is a general regulator of apoptosis (Ashkenazi et al. 2017). Only one of the novel TIR domain containing genes previously identified in actiniarians (Poole and Weis 2014; van der Burg et al. 2016) was found differentially expressed in multiple timepoints here, which was hypothetical gene with a predicted TIR-2 domain and with a possible novel domain. Whether this gene plays a significant role in regeneration is unknown.

Immune genes previously reported by Stewart *et al.* (2017) to be differentially expressed in *C. polypus* were not found to be differentially expressed in *E. pallida* and include: Alpha-2-macroglobulin (*A2M*), PF04505 domain (Interferon-induced transmembrane protein), Autophagy-related protein LC3C, and Fibrocystin-like gene (*PKHDL1*). The domain ‘Scavenger receptor cysteine rich’ (PF00530) is found in two differentially expressed genes (both *DMBT1*) [Supplementary Tables S5]. Interleukin-1 like receptor genes are differentially expressed in *C. polypus* – one putative interleukin-1 like receptor is identified here, and although it is annotated in the genome as ‘Toll-like receptor 2 type-2’ (gene1475), the predicted Pfam domains for the gene are typical of interleukin receptors. The lack of similarities between the two species could perhaps be explained by both the large scope of genes considered immune genes and the generally low expression of the immune system during regeneration.

The literature generally points towards an inverse relationship between the immune system and the regeneration response (Godwin and Brockes 2006; Altincicek and Vilcinskas 2008; Eming et al. 2009; Fukazawa et al. 2009; Peiris et al. 2014; van de Water et al. 2015; Abnave and Ghigo 2019). *Exaiptasia pallida* does lack some key immune genes (TLR and NF-κB), but the number of other immune genes (e.g., TIR-domain-containing genes and components of the complement system) are on par with other anthozoans (Poole and Weis 2014; Baumgarten et al. 2015; van der Burg et al. 2016) and notably, it has also been shown in *Hydra* that the lack of a TLR gene does not impede TLR pathway signalling (the canonical TLR protein is achieved through the scaffolding of two proteins) (Bosch et al. 2009). Whether or not this loss of some immune genes is part of a trade-off for increased regenerative ability is unknown, although the expression of immune genes in only the first three timepoints up to 8 hours suggests the immune response is very tightly controlled during regeneration. It should also be noted that the absence of these genes does not currently appear to reduce the ability of *E. pallida* to respond to bacterial treatment (Brown and Rodriguez-Lanetty 2015; Roesel and Vollmer 2019).

*Wnt* and *Wnt*-pathway related genes have been implicated in regeneration for many species (Chera et al. 2009; Duffy et al. 2010; Fumagalli et al. 2018; Hobmayer et al. 2000; Liu et al. 2013; Schaffer et al. 2016). In particular, the polar expression of Wnt appears to denote the regeneration of either head or body. Studies in *Hydra, Hydractinia* and *Nematostella* show that Wnt promotes head regeneration and suppresses stolon (or body) regeneration (Chera et al. 2009; Duffy et al. 2010; Hobmayer et al. 2000; Schaffer et al. 2016). The opposite is true in planarians; a study showed that head regeneration could be restored in the species *Dendrocoelum lacteum* by knocking down β-catenin in tail pieces and activation of β-catenin was “necessary and sufficient” to specify the tail axis (Liu et al. 2013). Multiple components of the Wnt pathway were expressed in *E. pallida*, (Figure 4) and all were upregulated which is consistent with data for other cnidarians. While it is interesting that this polarity appears to be somewhat ontologically reversed in planarians versus cnidarians (Schaffer et al. 2016), the recruitment of the same genes and gene pathways appears to be highly conserved regardless of the specific role in determining axis polarity.

A limitation of using software to model genes in a genome is that ORFs can be annotated that are not true ORFs, i.e., do not encode an expressed protein. Because of this, detecting novel, species-specific genes becomes more difficult as genes that appear as non-orthologous may be either truly novel or may be from incorrect gene models. To overcome this limitation, all 59 species-specific novels detected by OrthoFinder were used as a BLAST input against two independently assembled transcriptomes. Although these transcriptomes may in themselves not represent the entirety of expressed genes from *E. pallida*, they were at least able to be used to confidently identify 10 of the 59 genes as species-specific to *E. pallida.* The lack of Pfam or other annotations for these genes does not allow much to be deduced about their specific function, other than that they are potentially quite important to the regeneration response, as many of these species-specific novels are found throughout each timepoint (Supplementary Table S8.e) both up-and down-regulated. One of these species-specific genes identified was annotated as a collagen. Collagens are a component of the extracellular matrix and in sea anemones are an important part of the mesoglea, which is a connective tissue that allows the anemones to stretch and contract (Singer 1974; Shick 2012). An increase in the collagen present in the mesoglea has previously been identified in a morphological description of regeneration in *E. pallida* (synonymous with *Aiptasia diaphana*) (Singer 1974). The function of these collagens in regeneration have not been determined, but perhaps the increase in collagen expression could be related to the increase in rapid contractions and movement or ‘pulsing’ motions observed in *E. pallida* and in other regenerating cnidarians (Abrams et al. 2015; Stewart et al. 2017). Some of these species-specific genes may not be ‘novel’, since as more genomes are sequenced the genes may be shown to be present, but it is interesting that such a high number are not found in any of the closely related species included here. Regardless, there is an observable positive trend between the use of actiniarian and species-specific genes and regeneration, as exemplified by Figure 6.

As a final note, it is very interesting that no differential expression was observed in genes from *Symbiodinium minutum.* While only a small percentage of the reads aligned to the genome (∼8% aligned of ∼20 million reads) there was still a considerable number of genes that received at least one read mapping back. It is difficult to comment on why there was no observed differential expression, it could possibly be simply due to a lack of sequencing depth. A recent study that analysed differential expression in symbiotic and aposymbiotic *Exaiptasia* under bacterial immune challenge observed that there was no strong interaction between symbiotic state and bacterial exposure (Roesel and Vollmer 2019) and so perhaps the transcriptional regeneration response in *Exaiptasia* follows a fairly strict host-driven trajectory regardless of symbiotic state. In the future, testing the differences in regeneration response in both symbiotic and aposymbiotic *Exaiptasia* could provide some interesting clarification on this relationship.

## Conclusion

Altogether, this study illustrates that regeneration in *E. pallida* is characterised by a high degree of tissue plasticity, consisting of the mass movement of cells, coupled with a highly coordinated signalling response and cellular communication. Based on these transcriptional results, it can be hypothesised that *E pallida* will likely behave similarly to the model sea anemone *Nematostella* if cell proliferation were blocked, i.e., regeneration of *E. pallida* would proceed at first with complete wound healing, but tentacle elongation and pharynx formation would not occur (Amiel et al. 2015). The immune system is present during early regeneration (up to 8 hours) but does not dominate any of the enriched or depleted activities detected. Many species-specific genes were identified during regeneration in *E. pallida* with currently no known annotation or function. These genes, together with the high number of actiniarian-specific genes provide evidence that many genes differentially expressed during regeneration are highly taxonomically-restricted and likely allow *E. pallida* the ability to regenerate in a unique and rapid manner. Future studies will elucidate the specific role of these undescribed genes and further answer how, from a cellular perspective, *E. pallida* regenerates from either axes.

## Acknowledgements

The authors would like to acknowledge the Institute of Health and Biomedical Innovation (IHBI) for providing funding for the generation of raw reads for this project. Computational resources used in this work were provided by the HPC (High Performance Computing). Lab space was provided by MGRF (Molecular Genomics Research Facility) and technical support was provided by Vincent Chand and Sahana Manoli at MGRF at Queensland University of Technology, Brisbane, Australia. The authors would like to thank the members of the Prentis lab group, in particular Jessica O’Callaghan, for their insights and support. The authors would also like to thank the marine lab crew at QUT for their continual help with care and maintenance of the marine animals. Special thanks goes to Dr Libby Liggins (Massey University, New Zealand) for providing the *D. lineata* sample.

## Authors’ Contributions

C.V.D.B., A.P., J.S. and P.P. conceived and designed the research; C.V.D.B., J.S and H.S. performed the experiments; C.V.D.B and H.S. sequenced and assembled transcriptomes, C.V.D.B analysed the data; C.V.D.B., A.P., E.G., E.P., J.S., T.W. and P.P. interpreted results of experiments; C.V.D.B. prepared figures; C.V.D.B. and P.P. drafted the manuscript, C.V.D.B., A.P., E.G., E.P., J.S., H.S. T.W. and P.P. edited, revised and approved the final version of the manuscript.

## Compliance with Ethical Standards

This project did not require animal ethics approval, however, sample collection was authorised under the Fisheries Act 1994 (General Fisheries Permit), permit number: 166312

## Data availability statement

Raw reads for the *Exaiptasia pallida* regeneration time course are available on the Sequence Read Archive (SRA, NCBI) under BioProject accession number PRJNA507308. Full list of accession numbers are in Supplementary Table S1. Select gene annotation data is available in the supplementary data; additional gene annotations and read mapping files can be provided upon request. Raw reads for the three sea anemone transcriptomes generated here (*Diadumene lineata, Stichodactyla mertensii* and *Triactis producta*) can be found under BioProject accession number PRJNA507679.

## References

Abnave P, Ghigo E (2019) Role of the immune system in regeneration and its dynamic interplay with adult stem cells. Seminars in Cell & Developmental Biology 87:160– 168. https://doi.org/10.1016/j.semcdb.2018.04.002

Abrams MJ, Basinger T, Yuan W, et al (2015) Self-repairing symmetry in jellyfish through mechanically driven reorganization. PNAS 112:E3365–E3373. https://doi.org/10.1073/pnas.1502497112

Albertin CB, Simakov O, Mitros T, et al (2015) The octopus genome and the evolution of cephalopod neural and morphological novelties. Nature 524:220–224. https://doi.org/10.1038/nature14668

Altincicek B, Vilcinskas A (2008) Comparative analysis of septic injury-inducible genes in phylogenetically distant model organisms of regeneration and stem cell research, the planarian Schmidtea mediterranea and the cnidarian Hydra vulgaris. Front Zool 5:6. https://doi.org/10.1186/1742-9994-5-6

Alvarado AS, Tsonis PA (2006) Bridging the regeneration gap: genetic insights from diverse animal models. Nat Rev Genet 7:873–884. https://doi.org/10.1038/nrg1923

Amiel AR, Johnston HT, Nedoncelle K, et al (2015) Characterization of Morphological and Cellular Events Underlying Oral Regeneration in the Sea Anemone, Nematostella vectensis. Int J Mol Sci 16:28449–28471. https://doi.org/10.3390/ijms161226100

Ashkenazi A, Fairbrother WJ, Leverson JD, Souers AJ (2017) From basic apoptosis discoveries to advanced selective BCL-2 family inhibitors. Nature Reviews Drug Discovery 16:273–284. https://doi.org/10.1038/nrd.2016.253

Babonis LS, Martindale MQ (2017) Phylogenetic evidence for the modular evolution of metazoan signalling pathways. Phil Trans R Soc B 372:20150477. https://doi.org/10.1098/rstb.2015.0477

Baumgarten S, Simakov O, Esherick LY, et al (2015) The genome of Aiptasia, a sea anemone model for coral symbiosis. PNAS 112:11893–11898. https://doi.org/10.1073/pnas.1513318112

Bhattacharya D, Agrawal S, Aranda M, et al (2016) Comparative genomics explains the evolutionary success of reef-forming corals. eLife 5:e13288. https://doi.org/10.7554/eLife.13288

Bolger AM, Lohse M, Usadel B (2014) Trimmomatic: a flexible trimmer for Illumina sequence data. Bioinformatics 30:2114–2120. https://doi.org/10.1093/bioinformatics/btu170

Bosch TCG (2007) Why polyps regenerate and we don’t: Towards a cellular and molecular framework for Hydra regeneration. Dev Biol 303:421–433. https://doi.org/10.1016/j.ydbio.2006.12.012

Bosch TCG, Augustin R, Anton-Erxleben F, et al (2009) Uncovering the evolutionary history of innate immunity: the simple metazoan Hydra uses epithelial cells for host defence. Dev Comp Immunol 33:559–569. https://doi.org/10.1016/j.dci.2008.10.004

Brekhman V, Malik A, Haas B, et al (2015) Transcriptome profiling of the dynamic life cycle of the scypohozoan jellyfish Aurelia aurita. BMC Genomics 16:74. https://doi.org/10.1186/s12864-015-1320-z

Brockes JP, Kumar A (2008) Comparative Aspects of Animal Regeneration. Annu Rev Cell Dev Biol 24:525–549. https://doi.org/10.1146/annurev.cellbio.24.110707.175336

Brockes JP, Kumar A, Velloso CP (2001) Regeneration as an evolutionary variable. J Anat 199:3–11. https://doi.org/10.1046/j.1469-7580.2001.19910003.x

Brown T, Rodriguez-Lanetty M (2015) Defending against pathogens – immunological priming and its molecular basis in a sea anemone, cnidarian. Sci Rep 5:17425. https://doi.org/10.1038/srep17425

Browne EN (1909) The production of new hydranths in Hydra by the insertion of small grafts. J Exp Zool 7:1–23. https://doi.org/10.1002/jez.1400070102

Bucher M, Wolfowicz I, Voss PA, et al (2016) Development and Symbiosis Establishment in the Cnidarian Endosymbiosis Model Aiptasia sp. Sci Rep 6:19867. https://doi.org/10.1038/srep19867

Buzgariu W, Wenger Y, Tcaciuc N, et al (2018) Impact of cycling cells and cell cycle regulation on Hydra regeneration. Developmental Biology 433:240–253. https://doi.org/10.1016/j.ydbio.2017.11.003

Chera S, Ghila L, Dobretz K, et al (2009) Apoptotic Cells Provide an Unexpected Source of Wnt3 Signaling to Drive Hydra Head Regeneration. Dev Cell 17:279–289. https://doi.org/10.1016/j.devcel.2009.07.014

Clayton, Jr. WS (1985) Pedal laceration by the anemone Aiptasia pallida. Mar Ecol Prog Ser 21:75–80

DuBuc TQ, Traylor-Knowles N, Martindale MQ (2014) Initiating a regenerative response; cellular and molecular features of wound healing in the cnidarian Nematostella vectensis. BMC Biol 12:24. https://doi.org/10.1186/1741-7007-12-24

Duffy DJ, Plickert G, Kuenzel T, et al (2010) Wnt signaling promotes oral but suppresses aboral structures in Hydractinia metamorphosis and regeneration. Development 137:3057–3066. https://doi.org/10.1242/dev.046631

Eming SA, Hammerschmidt M, Krieg T, Roers A (2009) Interrelation of immunity and tissue repair or regeneration. Semin Cell Dev Biol 20:517–527. https://doi.org/10.1016/j.semcdb.2009.04.009

Forêt S, Knack B, Houliston E, et al (2010) New tricks with old genes: the genetic bases of novel cnidarian traits. Trends Genet 26:154–158. https://doi.org/10.1016/j.tig.2010.01.003

Fukazawa T, Naora Y, Kunieda T, Kubo T (2009) Suppression of the immune response potentiates tadpole tail regeneration during the refractory period. Development 136:2323–2327. https://doi.org/10.1242/dev.033985

Fumagalli MR, Zapperi S, Porta CAML (2018) Regeneration in distantly related species: common strategies and pathways. NPJ Syst Biol Appl 4:5. https://doi.org/10.1038/s41540-017-0042-z

Garza-Garcia AA, Driscoll PC, Brockes JP (2010) Evidence for the Local Evolution of Mechanisms Underlying Limb Regeneration in Salamanders. Integr Comp Biol 50:528–535. https://doi.org/10.1093/icb/icq022

Gierer A, Berking S, Bode H, et al (1972) Regeneration of Hydra from Reaggregated Cells. Nature New Biology 239:98–101. https://doi.org/10.1038/10.1038/newbio239098a0

Godwin JW, Brockes JP (2006) Regeneration, tissue injury and the immune response. J Anat 209:423–432. https://doi.org/10.1111/j.1469-7580.2006.00626.x

Grajales A, Rodríguez E (2016) Elucidating the evolutionary relationships of the Aiptasiidae, a widespread cnidarian–dinoflagellate model system (Cnidaria: Anthozoa: Actiniaria: Metridioidea). Mol Phylogenet Evol 94:252–263. https://doi.org/10.1016/j.ympev.2015.09.004

Grawunder D, Hambleton EA, Bucher M, et al (2015) Induction of Gametogenesis in the Cnidarian Endosymbiosis Model Aiptasia sp. Sci Rep 5:15677. https://doi.org/10.1038/srep15677

Gurtner GC, Werner S, Barrandon Y, Longaker MT (2008) Wound repair and regeneration. Nature 453:314–321. https://doi.org/10.1038/nature07039

Haas BJ, Papanicolaou A, Yassour M, et al (2013) De novo transcript sequence reconstruction from RNA-seq using the Trinity platform for reference generation and analysis. Nat Protoc 8:1494–1512. https://doi.org/10.1038/nprot.2013.084

Hobmayer B, Rentzsch F, Kuhn K, et al (2000) WNT signalling molecules act in axis formation in the diploblastic metazoan Hydra. Nature 407:186–189. https://doi.org/10.1038/35025063

Holstein T w., Hobmayer E, Technau U (2003) Cnidarians: An evolutionarily conserved model system for regeneration? Dev Dyn 226:257–267. https://doi.org/10.1002/dvdy.10227

Huang C, Morlighem J-ÉR, Zhou H, et al (2016) The Transcriptome of the Zoanthid Protopalythoa variabilis (Cnidaria, Anthozoa) Predicts a Basal Repertoire of Toxin-like and Venom-Auxiliary Polypeptides. Genome Biol Evol 8:3045–3064. https://doi.org/10.1093/gbe/evw204

Liu S-Y, Selck C, Friedrich B, et al (2013) Reactivating head regrowth in a regeneration-deficient planarian species. Nature 500:81–84. https://doi.org/10.1038/nature12414

Manuel M (2009) Early evolution of symmetry and polarity in metazoan body plans. Comptes Rendus Biologies 332:184–209. https://doi.org/10.1016/j.crvi.2008.07.009

Miljkovic-Licina M, Chera S, Ghila L, Galliot B (2007) Head regeneration in wild-type hydra requires de novo neurogenesis. Development 134:1191–1201. https://doi.org/10.1242/dev.02804

Miller DJ, Ball EE, Technau U (2005) Cnidarians and ancestral genetic complexity in the animal kingdom. Trends Genet 21:536–539. https://doi.org/10.1016/j.tig.2005.08.002

Morgan TH (1901) Regeneration in the Egg, Embryo, and Adult. The American Naturalist 35:949–973

Oakley CA, Ameismeier MF, Peng L, et al (2016) Symbiosis induces widespread changes in the proteome of the model cnidarian Aiptasia. Cell Microbiol 18:1009–1023. https://doi.org/10.1111/cmi.12564

Passamaneck YJ, Martindale MQ (2012) Cell proliferation is necessary for the regeneration of oral structures in the anthozoan cnidarian Nematostella vectensis. BMC Dev Biol 12:34. https://doi.org/10.1186/1471-213X-12-34

Peiris TH, Hoyer KK, Oviedo NJ (2014) Innate immune system and tissue regeneration in planarians: An area ripe for exploration. Semin Immunol 26:295–302. https://doi.org/10.1016/j.smim.2014.06.005

Petersen HO, Höger SK, Looso M, et al (2015) A comprehensive transcriptomic and proteomic analysis of Hydra head regeneration. Mol Biol Evol 32:1928–1947. https://doi.org/10.1093/molbev/msv079

Poole AZ, Kitchen SA, Weis VM (2016) The Role of Complement in Cnidarian-Dinoflagellate Symbiosis and Immune Challenge in the Sea Anemone Aiptasia pallida. Front Microbiol 7:. https://doi.org/10.3389/fmicb.2016.00519

Poole AZ, Weis VM (2014) TIR-domain-containing protein repertoire of nine anthozoan species reveals coral–specific expansions and uncharacterized proteins. Dev Comp Immunol 46:480–488. https://doi.org/10.1016/j.dci.2014.06.002

Poss KD (2010) Advances in understanding tissue regenerative capacity and mechanisms in animals. Nat Rev Genet 11:710–722. https://doi.org/10.1038/nrg2879

Putnam NH, Srivastava M, Hellsten U, et al (2007) Sea Anemone Genome Reveals Ancestral Eumetazoan Gene Repertoire and Genomic Organization. Science 317:86–94. https://doi.org/10.1126/science.1139158

Rodríguez E, Barbeitos MS, Brugler MR, et al (2014) Hidden among Sea Anemones: The First Comprehensive Phylogenetic Reconstruction of the Order Actiniaria (Cnidaria, Anthozoa, Hexacorallia) Reveals a Novel Group of Hexacorals. PLOS ONE 9:e96998. https://doi.org/10.1371/journal.pone.0096998

Roesel CL, Vollmer SV (2019) Differential gene expression analysis of symbiotic and aposymbiotic Exaiptasia anemones under immune challenge with Vibrio coralliilyticus. Ecology and Evolution 0: https://doi.org/10.1002/ece3.5403

Schaffer AA, Bazarsky M, Levy K, et al (2016) A transcriptional time-course analysis of oral vs. aboral whole-body regeneration in the Sea anemone Nematostella vectensis. BMC Genomics 17:718. https://doi.org/10.1186/s12864-016-3027-1

Shick JM (2012) A Functional Biology of Sea Anemones. Chapman & Hall, London

Shoguchi E, Shinzato C, Kawashima T, et al (2013) Draft assembly of the Symbiodinium minutum nuclear genome reveals dinoflagellate gene structure. Curr Biol 23:1399– 1408. https://doi.org/10.1016/j.cub.2013.05.062

Singer II (1974) An electron microscopic and autoradiographic study of mesogleal organization and collagen synthesis in the sea anemone Aiptasia diaphana. Cell Tissue Res 149:537–554. https://doi.org/10.1007/BF00223031

Singer II (1971) Tentacular and oral-disc regeneration in the sea anemone, Aiptasia diaphana III. J Embryo Exp Morph 26:253–270

Singer II, Palmer JD (1969) Tentacular and oral-disc regeneration in the sea anemone, Aiptasia diaphana II. Naturwissenschaften 56:574–575. https://doi.org/10.1007/BF00597293

Sinigaglia C, Busengdal H, Leclère L, et al (2013) The Bilaterian Head Patterning Gene six3/6 Controls Aboral Domain Development in a Cnidarian. PLOS Biol 11:e1001488. https://doi.org/10.1371/journal.pbio.1001488

Srivastava M, Begovic E, Chapman J, et al (2008) The *Trichoplax* genome and the nature of placozoans. Nature 454:955–960. https://doi.org/10.1038/nature07191

Srivastava M, Simakov O, Chapman J, et al (2010) The *Amphimedon queenslandica* genome and the evolution of animal complexity. Nature 466:720–726. https://doi.org/10.1038/nature09201

Stewart ZK, Pavasovic A, Hock DH, Prentis PJ (2017) Transcriptomic investigation of wound healing and regeneration in the cnidarian *Calliactis polypus*. Sci Rep 7:41458. https://doi.org/10.1038/srep41458

Surm JM, Smith HL, Madio B, et al (2019) A process of convergent amplification and tissue-specific expression dominates the evolution of toxin and toxin-like genes in sea anemones. Molecular Ecology 28:2272–2289. https://doi.org/10.1111/mec.15084

Tanaka EM, Reddien PW (2011) The Cellular Basis for Animal Regeneration. Dev Cell 21:172–185. https://doi.org/10.1016/j.devcel.2011.06.016

Tiozzo S, Copley RR (2015) Reconsidering regeneration in metazoans: an evo-devo approach. Front Ecol Evol 3:67. https://doi.org/10.3389/fevo.2015.00067

Trembley A (1744) Mémoires pour servir a l’histoire d’un genre de polypes d’eau douce, a bras en forme de cornes. J. and H. Verbeek

van de Water JAJM, Ainsworth TD, Leggat W, et al (2015) The coral immune response facilitates protection against microbes during tissue regeneration. Mol Ecol 24:3390– 3404. https://doi.org/10.1111/mec.13257

van der Burg CA, Prentis PJ, Surm JM, Pavasovic A (2016) Insights into the innate immunome of actiniarians using a comparative genomic approach. BMC Genomics 17:850. https://doi.org/10.1186/s12864-016-3204-2

Wang X, Liew YJ, Li Y, et al (2017) Draft genomes of the corallimorpharians Amplexidiscus fenestrafer and Discosoma sp. Mol Ecol Resour 17:e187–e195. https://doi.org/10.1111/1755-0998.12680

Yum LK, Baumgarten S, Röthig T, et al (2017) Transcriptomes and expression profiling of deep-sea corals from the Red Sea provide insight into the biology of azooxanthellate corals. Sci Rep 7:6442. https://doi.org/10.1038/s41598-017-05572-x

Zhang G, Fang X, Guo X, et al (2012) The oyster genome reveals stress adaptation and complexity of shell formation. Nature 490:49–54. https://doi.org/10.1038/nature11413

