## Supplementary File 1 for "The rapid regenerative response of a model sea anemone species *Exaiptasia pallida* is characterised by tissue plasticity and highly coordinated cell communication"

### BioRxiv

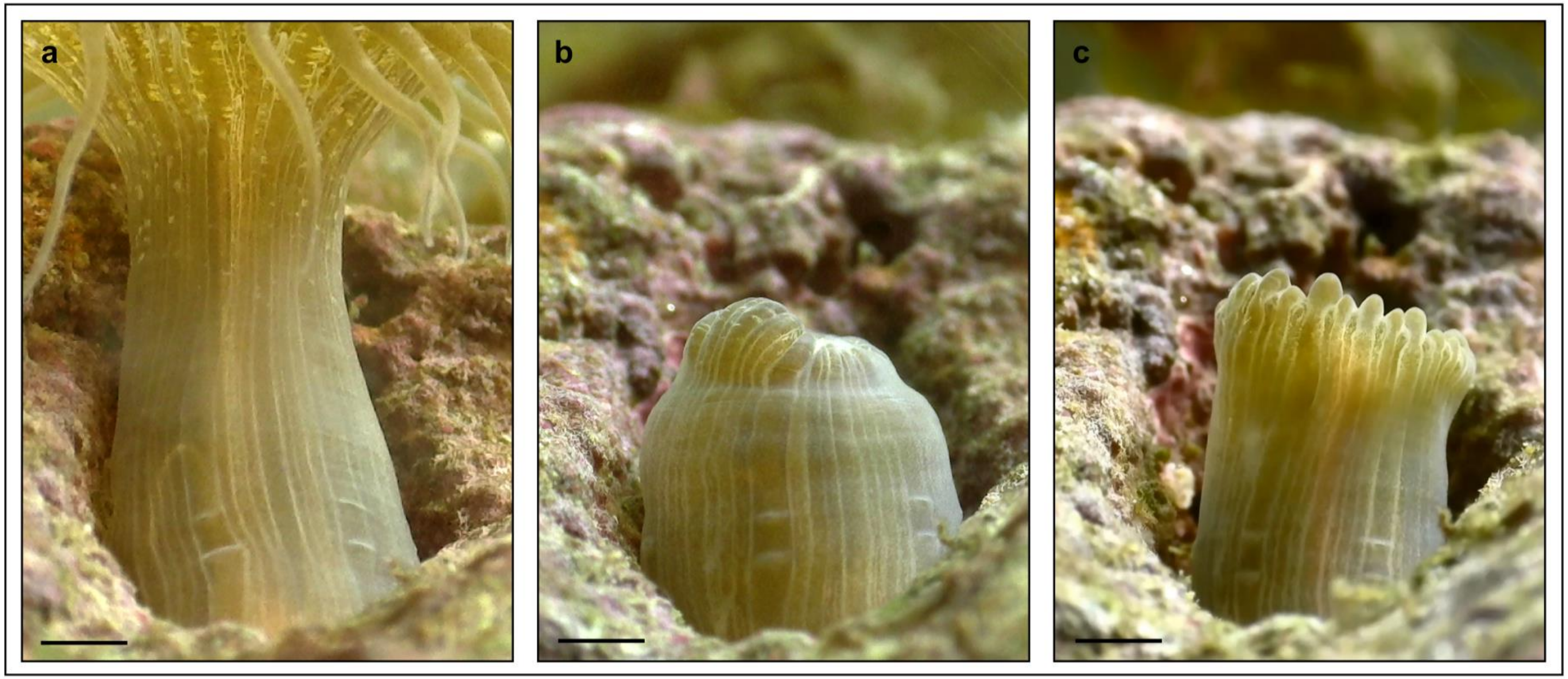


**Figure S1. Time series of tentacle regrowth shows no substantial body column growth occurs shortly following head amputation**. Image **a** shows the anemone immediately preceding head amputation. Image **b** shows the same anemone 24 hours p.h.a. Image **c** shows the same anemone four days p.h.a. The tentacles have for the most part grown directly out of the body column at the location where the wound has closed, without first growing the body column back to the original height before amputation. Scale = approx. 5 mm

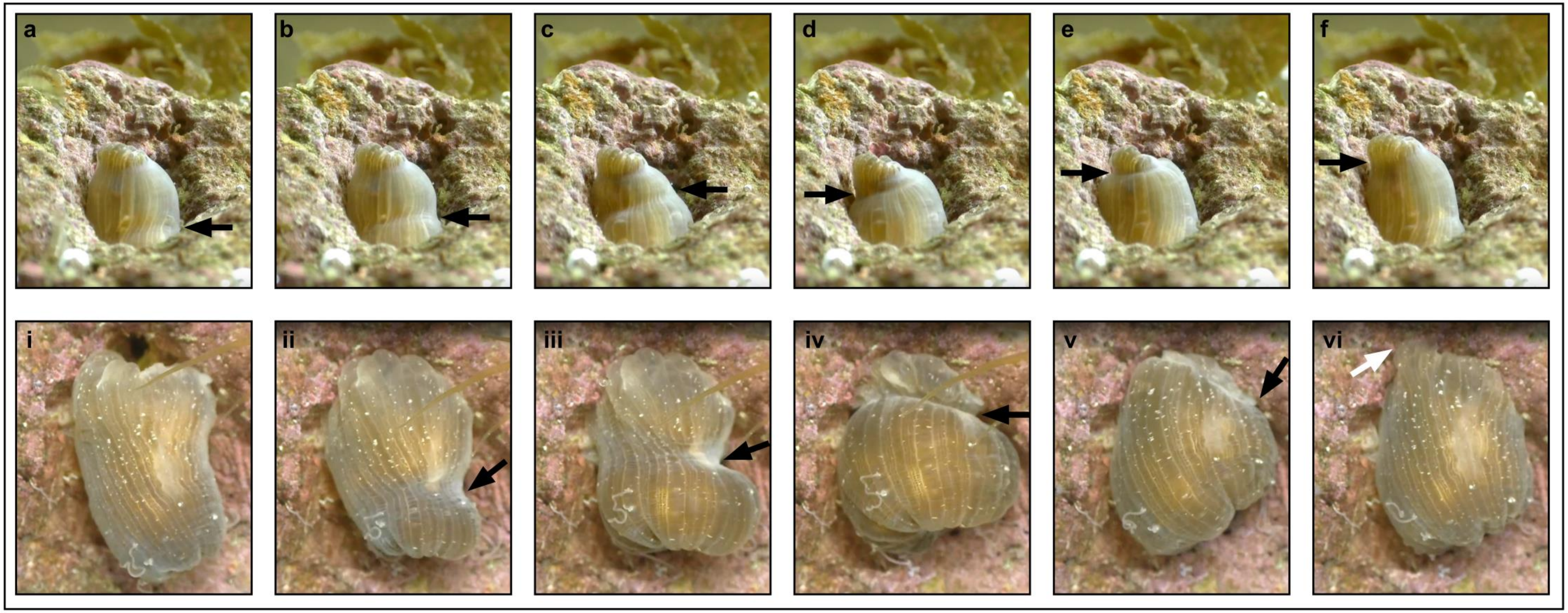


**Figure S2. Images of the 'upwards' and 'downwards' pulsing motion observed in *E. pallida* following head amputation**. Series **a-f** (top left to top right) show an upwards pulse; series **i-vi** (bottom left to bottom right) show a downwards pulse. The black arrows indicate the 'pinch' site, where the body contracts inwards. The white arrow on image **vi** indicates a portion of the foot that has become attached to the rock following a downwards pulse. Both series of images occur over about 2.5-3 minutes real-time (~1 second of film).
