## Supplementary File 2 for "The rapid regenerative response of a model sea anemone species *Exaiptasia pallida* is characterised by tissue plasticity and highly coordinated cell communication"

### BioRxiv

### Title

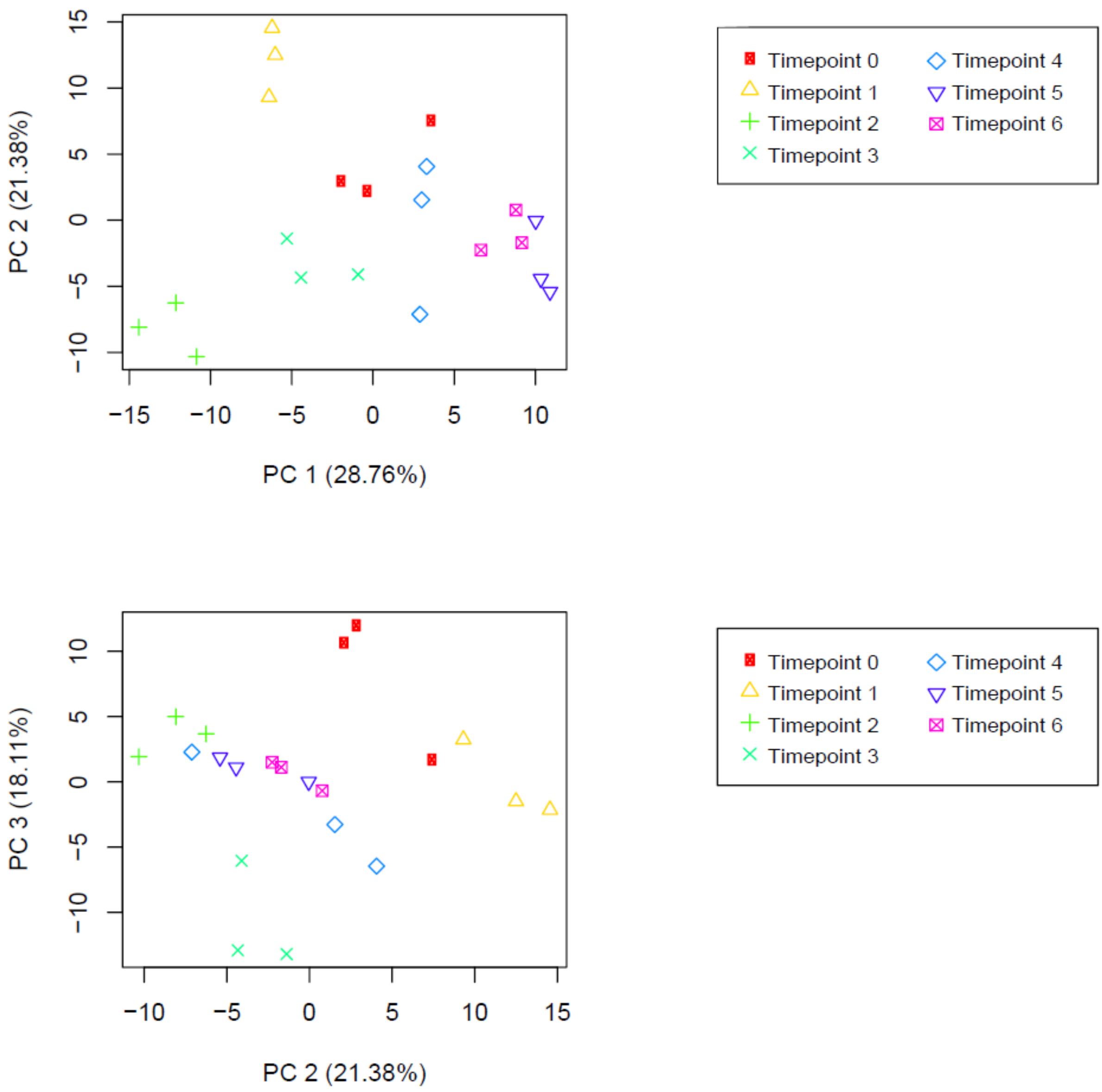


**Figure S3. Principle components analysis (PCA) plots of all 21 datasets**. Timepoint 0: datasets 1, 2, 3; timepoint 1: datasets 4, 5, 6; timepoint 2: datasets 7, 8, 9; timepoint 3: datasets 10, 11, 12; timepoint 4: datasets 13, 14, 15; timepoint 5: datasets 16, 17, 18; timepoint 6: datasets 19, 20, 21. Produced using statistically significant differentially expressed genes.

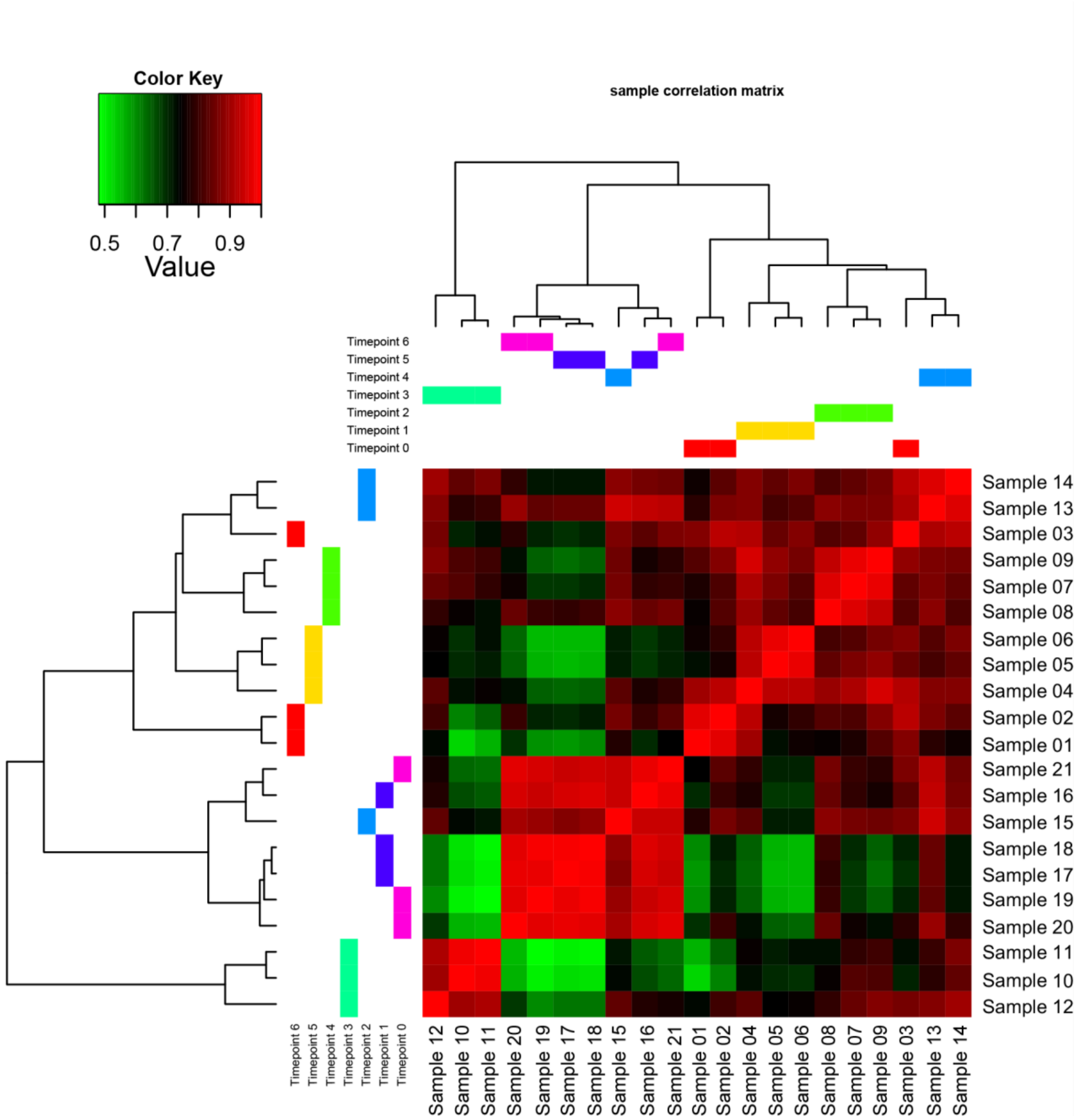


**Figure S4. Sample correlation matrix using statistically significant differentially expressed genes.**

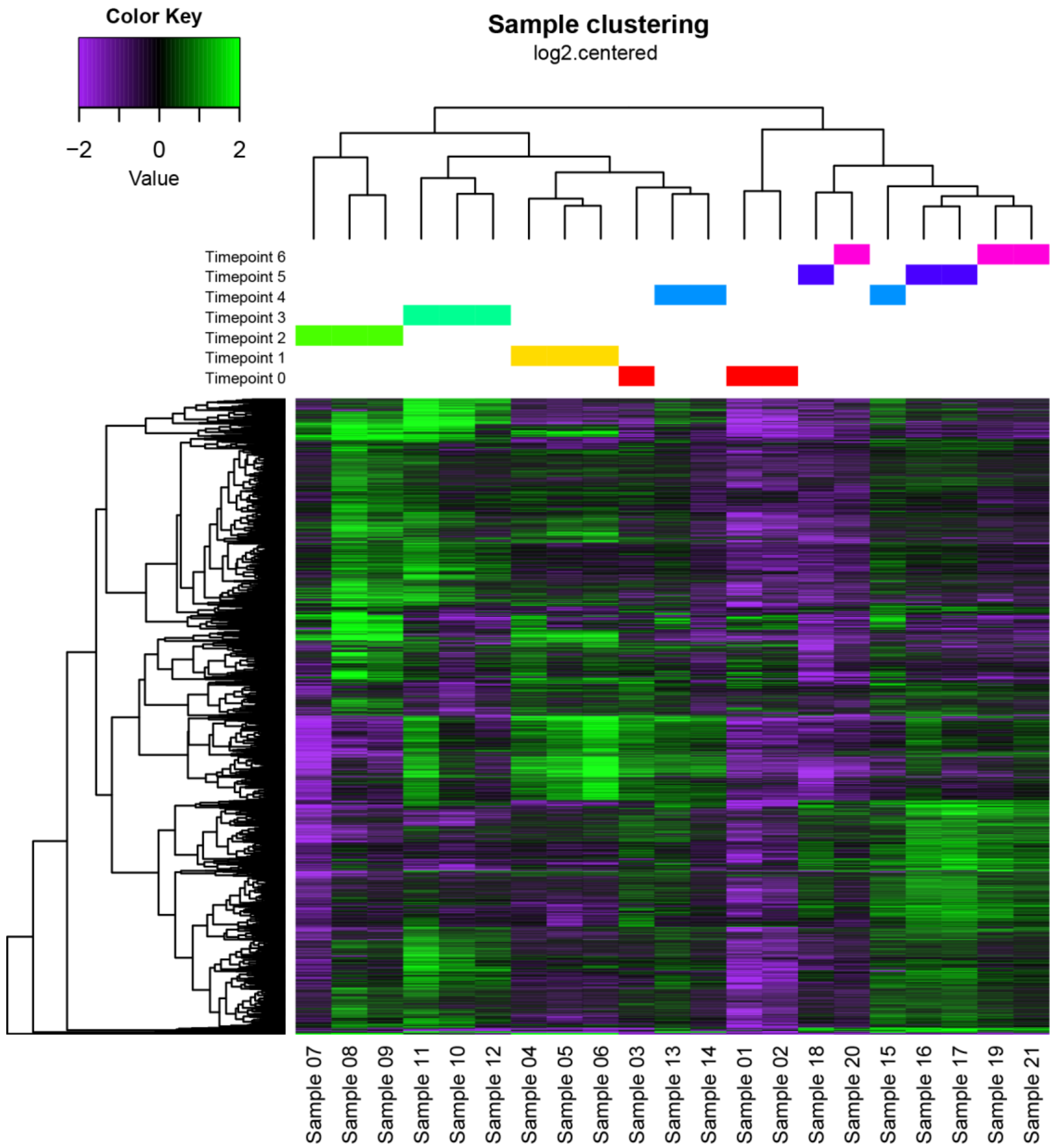


**Figure S5. Sample clustering heatmap using statistically significant differentially expressed genes.**

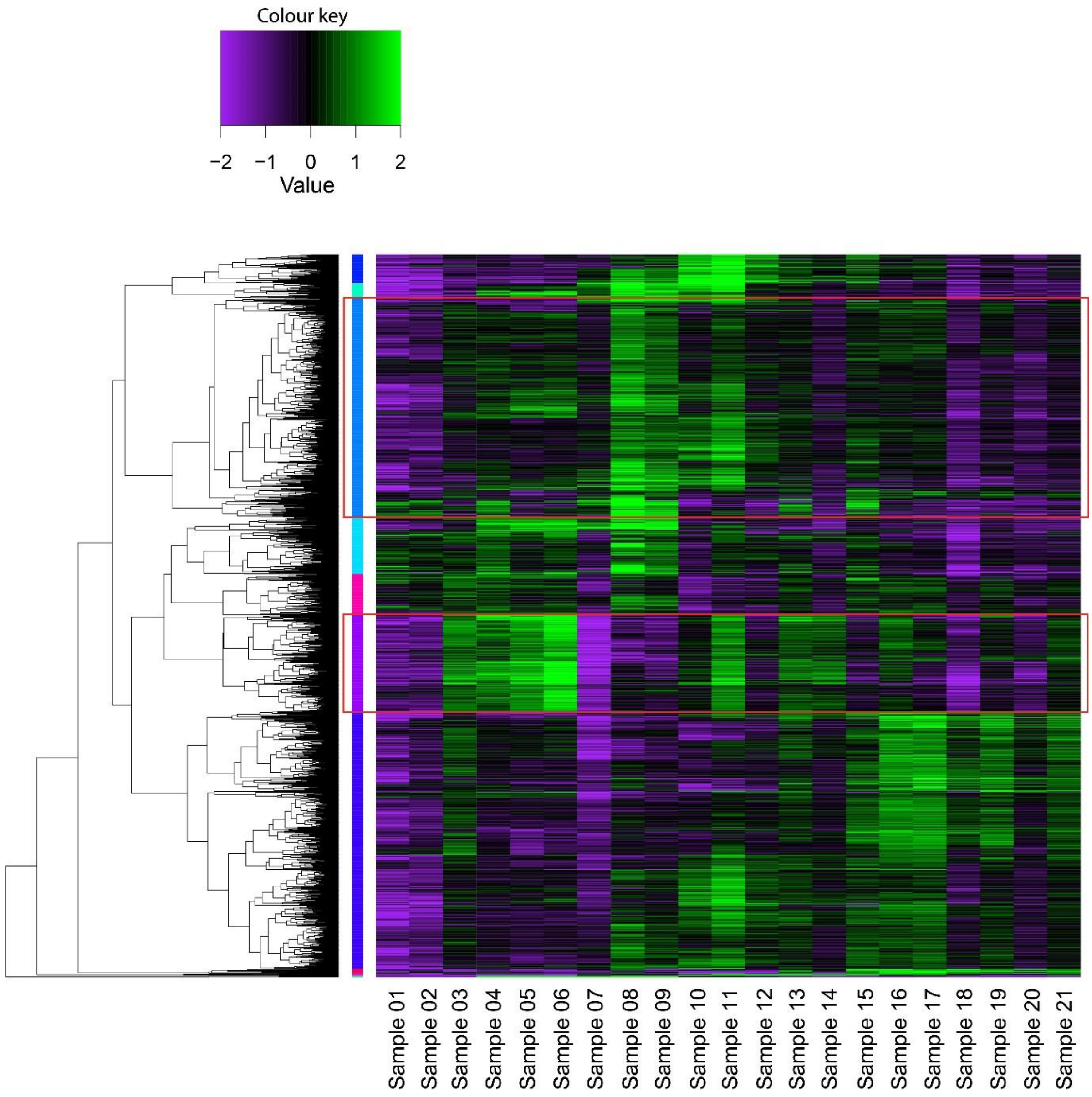


**Figure S6. Heatmap of statistically significant differentially expressed genes showing the subclusters recovered after cutting the tree at P50.** Subcluster 4 (blue band) and subcluster 7 (purple band) are highlighted with red boxes.

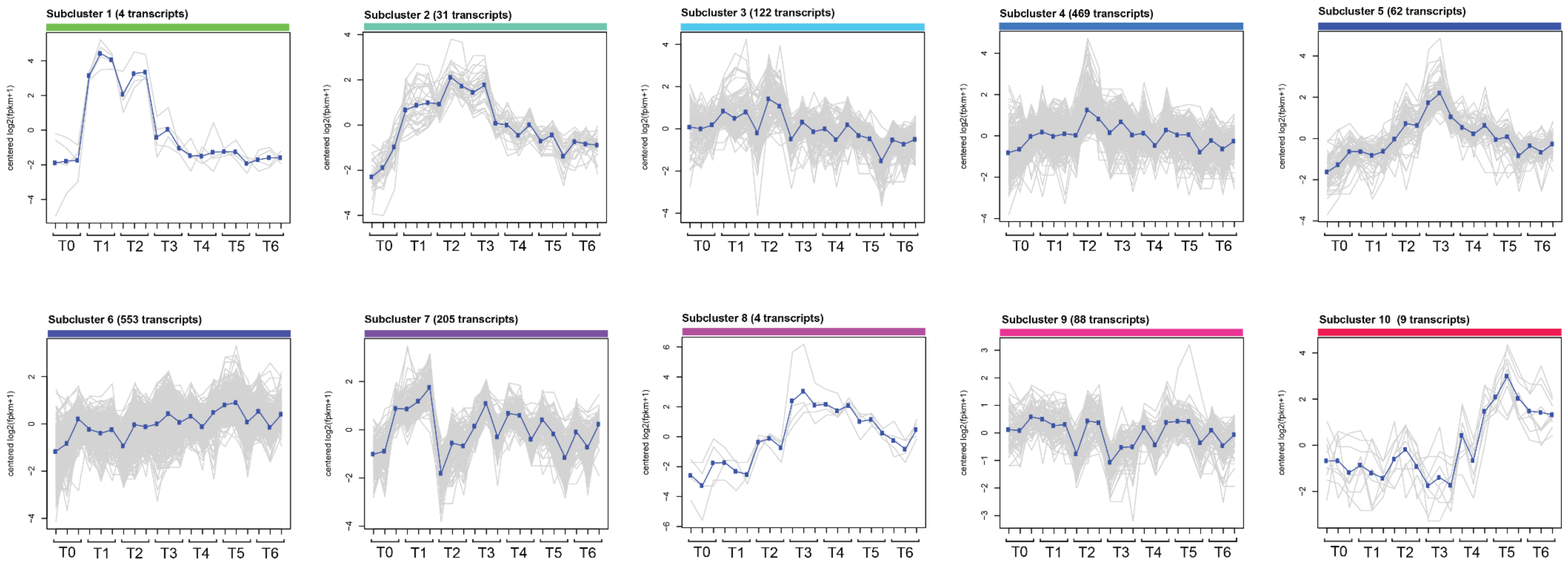


**Figure S7. Subclusters recovered from cutting the heatmap tree at 50% of max height.** Subclusters show the pattern of gene expression change over 7 timepoints, using centered log2 fold change as the metric for expression.
